# NgAgo DNA endonuclease activity enhances homologous recombination in *E. coli*

**DOI:** 10.1101/597237

**Authors:** Kok Zhi Lee, Michael A. Mechikoff, Archana Kikla, Arren Liu, Paula Pandolfi, Kevin Fitzgerald, Frederick S. Gimble, Kevin V. Solomon

**Affiliations:** Department of Agricultural and Biological Engineering, Purdue University, West Lafayette, IN 47906, USA; Department of Biological Sciences, Purdue University, West Lafayette, IN 47906, USA; Purdue University Interdisciplinary Life Science Program (PULSe), Purdue University, West Lafayette, IN 47906, USA; Department of Biochemistry, Purdue University, West Lafayette, IN, 47906, USA

## Abstract

Prokaryotic Argonautes (pAgos) have been proposed as more flexible tools for gene-editing as they do not require sequence motifs adjacent to their targets for function, unlike popular CRISPR/Cas systems. One promising pAgo candidate, from the halophilic archaeon *Natronobacterium gregoryi* (NgAgo), however, has been the subject of intense debate regarding its potential in eukaryotic systems. Here, we revisit this enzyme and characterize its function in prokaryotes. NgAgo expresses poorly in non-halophilic hosts with the majority of protein being insoluble and inactive even after refolding. However, we report that the soluble fraction does indeed act as a DNA endonuclease. Structural homology modelling revealed that NgAgo shares canonical domains with other catalytically active pAgos but also contains a previously unrecognized single-stranded DNA binding domain (repA). Both repA and the canonical PIWI domains participate in DNA cleavage activities of NgAgo. We showed that NgAgo can be programmed with guides to cleave specific DNA *in vitro* and in *E.coli*. We also found that these endonuclease activities are essential for enhanced NgAgo-guided homologous recombination, or gene-editing, in *E. coli*. Collectively, our results demonstrate the potential of NgAgo for gene-editing and reconciles seemingly contradictory reports.

---

Long pAgos are programmable endonucleases that bind single-stranded DNA and/or RNA molecules as guides, which then prime the enzyme for nicking of complementary target DNA, RNA, or both^1^. Double stranded DNA cleavage requires two complementary guides. DNA cleavage induced by pAgos enables DNA repair and editing, potentially forming an alternative gene editing platform to standard CRISPR-based tools. Unlike Cas9-based gene editing strategies, however, pAgos have the distinct advantage of not requiring a protospacer adjacent motif (PAM) for function^2–5^. Thus, pAgos are not limited to targets flanked by PAM sites and can potentially cut any DNA target regardless of composition. Despite this potential, no pAgo has been developed that rivals the simplicity and function of Cas9-based strategies.

Target recognition and cleavage is enabled by four canonical domains^3^: N (N-terminal), PAZ (PIWI-Argonaute-Zwille), MID (middle), and PIWI (P element-induced wimpy testis) domains. The N-terminal domain is essential for target cleavage^6,7^ and dissociation of cleaved strands^7,8^, although the detailed mechanism remains poorly understood. The MID domain interacts with the 5’-end of the guide^9^ and promotes binding to its target^10^. The PAZ domain interacts with the 3’ end of the guide^11–14^, protecting it from degradation^15^. Finally, the PIWI domain plays a pivotal role in nucleic acid cleavage via the conserved catalytic tetrad, DEDX (D: aspartate, E: glutamate, X: histidine, aspartate or asparagine)^16^.

Recent emerging evidence also suggests a role for accessory proteins in pAgo activity. Within prokaryote genomes, pAgos are often organized in operons with ssDNA binding proteins and helicases among other DNA modifying proteins^17^ hinting at concerted function *in vivo*. Supplementing a pAgo with these proteins *in vitro* enhances reaction rates and target specificity, reduces biases in substrate composition preferences, and enables activity on more topologically diverse substrates^18^. These effects are observed with several homologs of these accessory proteins for multiple pAgos. Moreover, pAgos also copurify with helicases, ssDNA binding proteins, and recombinases from both native and heterologous hosts^19,20^ indicating conserved physical interactions in different prokaryotes. Given the need for these and potentially other unrecognized accessory proteins, *in vivo* evaluation of pAgos may more accurately reflect their activity.

Despite the potential for programmable cleavage activities by long pAgos, currently characterized pAgos including TtAgo^2^, MpAgo^5^, PfAgo^21^ and MjAgo^3,22^ work at very high temperatures (>55 °C), making them infeasible for gene editing and *in vivo* testing in common mesophilic organisms. The halophilic Argonaute from the archaeon *Natronobacterium gregoryi* (NgAgo) was recently put forth as a promising candidate for pAgo-mediated gene editing, as it was believed to be active at mesophilic (~37°C) temperatures^23^. However, these claims have since been refuted due to an inability to demonstrate *in vitro* DNA cleavage or to replicate these findings in a number of eukaryotic hosts ^24–28^. NgAgo expression is poor, presumably due to its halophilic characteristics that make low salt expression challenging^29,30^. Thus, all published *in vitro* cleavage assays have relied on refolded protein^18,31^, which may be non-functional, resulting in the inconclusive results. Nonetheless, recent work by Fu and colleagues demonstrated that NgAgo may still have potential as a gene editor for prokaryotic hosts^19^. While the authors were able to confirm that gene-editing was mediated by homologous recombination via RecA^19^, which physically associated with NgAgo in an unanticipated manner, the specific role of NgAgo remained unclear. Here, we demonstrate that NgAgo is indeed a DNA endonuclease by identifying residues that are required for DNA cleavage, and we provide evidence that this activity is essential for NgAgo-mediated gene editing via homologous recombination repair.

## RESULTS

### NgAgo has canonical N-terminal, PIWI, MID, and PAZ domains, and a putative single stranded DNA binding (repA) domain

Given the ongoing debate of the function of NgAgo, we analyzed its sequence (IMG/M Gene ID: 2510572918) with Phyre 2^32^ and HHpred^33,34^ to predict its structure based on characterized structural homologs. Phyre 2 and HHpred analyses found with high confidence (probability = 100%) that NgAgo shares structural features with catalytically active pAgos and eukaryotic Agos (eAgos) including archaeal MjAgo, bacterial TtAgo, and eukaryotic hAgo2 (Supplementary Table 1 and 2). Since MjAgo is the only characterized pAgo from Archaea, we used it as a template for comparative modelling. The predicted NgAgo structure is similar to the crystal structure of MjAgo, consisting of canonical N-terminal, PAZ, MID, and PIWI domains (Fig. 1a and b). However, the N-terminal domain of NgAgo, which plays a key role in targeted cleavage, is truncated, relative to MjAgo. This may suggest a novel mechanism for strand displacement and binding.

**Figure 1.**
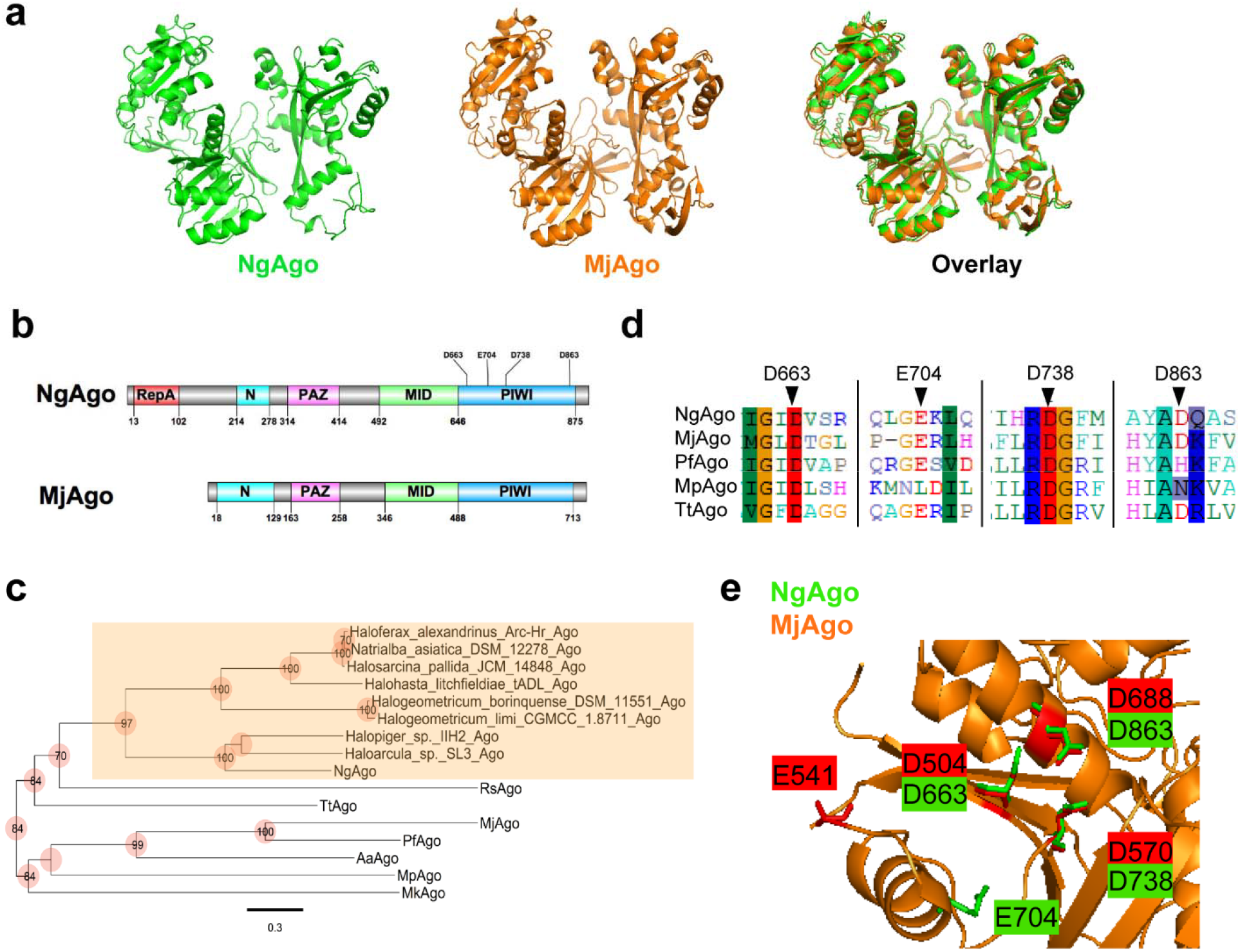
NgAgo belongs to a distinct clade of pAgos with a catalytic DEDX tetrad and novel repA domain. **a,** Phyre 2 simulation 3D structure based on MjAgo structure (PDB: 5G5T). NgAgo structure is similar to MjAgo structure except for at the N-terminal domain. **b**, Domain architecture analysis of NgAgo◻based on Phyre2 and HHpred reveals that NgAgo has an uncharacterized repA domain, a truncated N-terminal domain, a MID domain, and a PIWI domain. **c**, Phylogenetic analysis of repA-containing pAgos (orange shaded) found from BLASTP against all isolates via JGI-IMG portal and other characterized pAgos. **d**, The catalytic tetrad of NgAgo is conserved with catalytically active pAgos including MjAgo, PfAgo, MpAgo, and TtAgo in sequence alignment. **e**, All residues of the catalytic tetrad (D663, E704, D738, and D863) DEDD, except E704 are structurally colocalized with the catalytic tetrad of MjAgo (D504, E541, D570, and D688).

Structural analysis also identified an uncharacterized oligonucleotide/oligosaccharide-binding (OB) fold domain between residues 13-102 of NgAgo that commonly binds single-stranded DNA in eukaryotes and prokaryotes^35^ (Fig. 1b). This OB domain has recently been identified as a new feature of pAgos^17^. As repA proteins were the most common matches on both Phyre 2 and HHpred, we will refer to this OB domain as repA (Supplementary Tables 3 and 4). While the repA domain is absent in all characterized pAgos, at least 12 sequenced pAgo homologs share this domain. Phylogenetic analysis showed that all the repA-containing pAgos were from halophilic Archaea forming a clade that is distinct from that of the current well-characterized pAgos (Fig. 1c). This monophyletic group of repA-containing pAgos may represent a distinct class of pAgos that is currently unrecognized in the literature^17^. Moreover, its unique presence within halophiles may be evidence that the repA domain is required for function in high salt environments, potentially replacing the role of the canonical N-terminal domain, which was then truncated through evolution.

Our analysis of NgAgo also confirmed the presence of a conserved catalytic tetrad, DEDX (X: H, D or N)^16^, which is critical for nucleic acid cleavage by the PIWI domain of Argonautes. The catalytic tetrad (D663, E704, D738, and D863) of NgAgo aligns well with those from other catalytically active pAgos, including MjAgo^3^, PfAgo^21^, MpAgo^5^, and TtAgo^2^ (Fig. 1d). Moreover, structural alignment of NgAgo and MjAgo display good colocalization of D663, D738, and D863 within the catalytic tetrad suggesting that NgAgo may have similar nucleic acid cleavage activity (Fig. 1e).

### Soluble, but not refolded, NgAgo exhibits DNA cleavage activity *in vitro*

As halophilic proteins tend to be insoluble in low-salt environments due to their sequence adaptations^29,30,36^, we first optimized expression conditions to obtain more soluble NgAgo protein (Supplementary Fig. 1). NgAgo was still unstable in optimal expression conditions, as evidenced by truncated peptide products (Supplementary Fig. 1b). We purified wildtype NgAgo from both the soluble and insoluble fractions to test for 5’P-ssDNA guide-dependent DNA cleavage (Supplementary Fig. 2). Insoluble NgAgo was refolded during purification using established methods^31^. Purified NgAgo from the soluble fraction (sNgAgo) nicks plasmid DNA and genomic DNA, independent of a guide (Supplementary Fig. 3a), as evidenced by the presence of the nicked and linearized plasmid. However, refolded NgAgo from the insoluble lysate fraction (rNgAgo) has little or no activity on DNA (Supplementary Fig. 3b), consistent with a study by Ye and colleagues^31^.

### RepA and PIWI domains of NgAgo are required for DNA cleavage

To rule out the possibility of non-specific host nuclease impurities (Supplementary Fig. 4), we pursued cell-free expression of NgAgo. This approach has successfully been used to rapidly prototype other endonucleases including CRISPR-Cas endonuclease^37^. NgAgo expression was induced in the presence of 5’ phosphorylated guides that targeted a plasmid substrate, pNCS-mNeonGreen (Figs 2a,b). NaCl was supplemented after expression to promote proper folding of the halophilic enzyme (Fig. 2c, materials and methods). To identify regions critical for DNA cleavage, we constructed and expressed the repA domain of NgAgo (residues 1-102), a truncated NgAgo without the repA domain (residues 105-887, referred to as N-del) and D663A/D738A point mutations in the full-length protein and N-del variant (Fig. 2d). D663A/D738A is a double mutant within the catalytic tetrad that corresponds to the catalytic double mutant D478A/D546A of TtAgo^2^, which lost all cleavage activities^2,38^.

**Figure 2.**
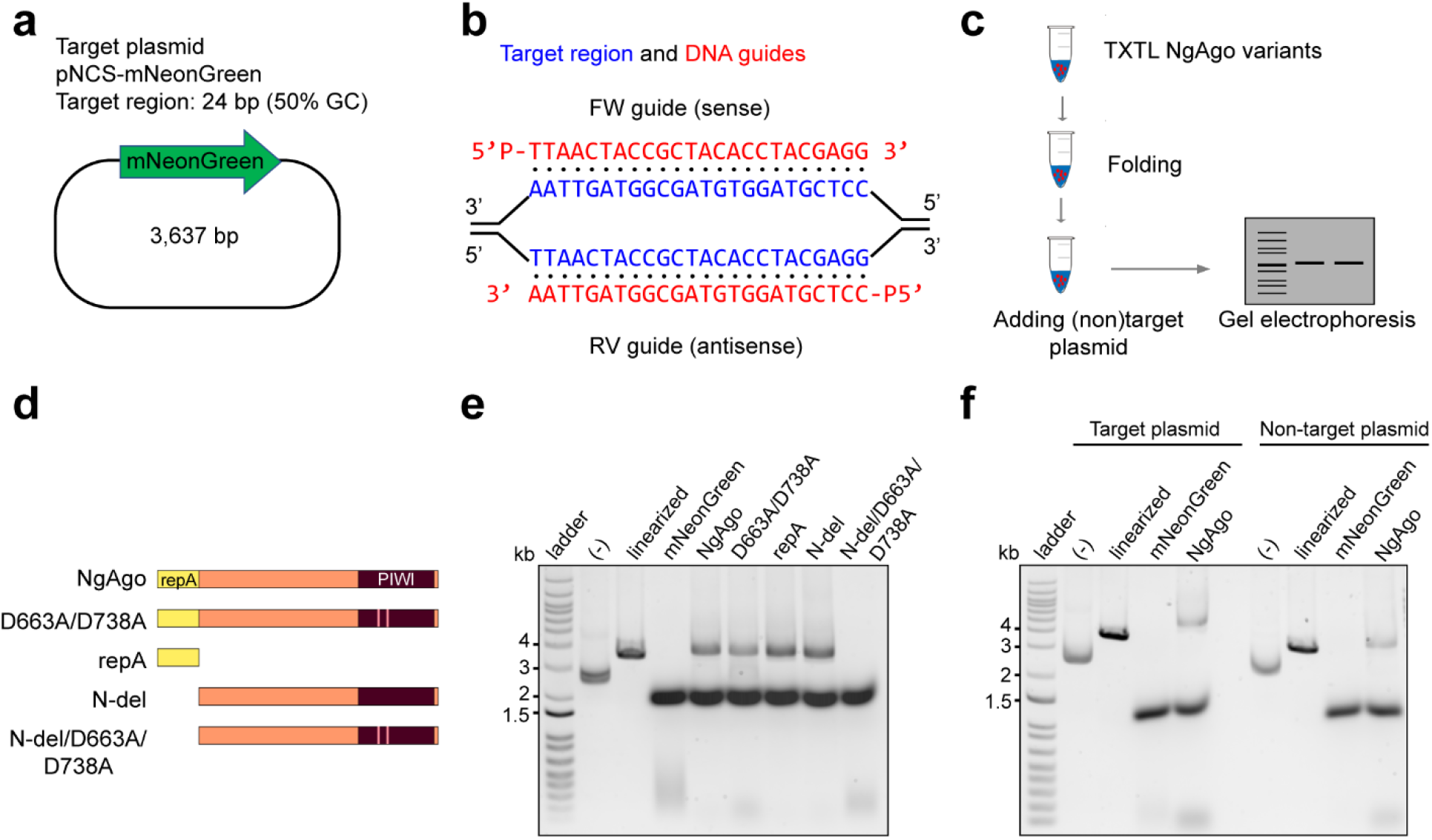
NgAgo variants degrade plasmid DNA *in vitro* via the repA domain and D663/D738 residues in the PIWI domain. **a**, Target plasmid pNCS-mNeonGreen contains a 24-base pair target site with 50% GC content. b, 5’ phosphorylated DNA guides binds to target sequence in pNCS-mNeonGreen. c, Procedure for bacterial cell-free-system production of NgAgo and DNA degradation assessment. d, NgAgo variants used in the *in vitro* assay to identify which domain is essential for nicking and cleaving activity. **e**, Plasmids were treated with NgAgo variants or mNeonGreen as a endonuclease negative control for an hour before analysis on an agarose gel. Wildtype and D663A/D738A degrades plasmids DNA while N-del degrades plasmid DNA with compromised activity. N-del/D663A/D738A loses the ability to degrade plasmid DNA. **f**, NgAgo degrades both target plasmid pNCS-mNeonGreen and non-target plasmid pBSI-SceI(E/H). Negative controls (-) are plasmids without any treatments.

Not all NgAgo variants displayed DNA cleavage activity, confirming that previously observed DNA cleavage could be attributed to NgAgo activity (Fig. 2e). Both wildtype NgAgo and D663A/D738A linearized substrate DNA suggesting catalytic activity beyond the PIWI domain^31^ or rescue of functionality by other domains even in the presence of a PIWI mutation. Both repA and PIWI domains participate in DNA cleavage and with each being sufficient for activity as cleavage was retained in both repA and N-del mutants. While it is unclear how the repA domain might lead to DNA damage, its single-stranded DNA binding activity in isolation may be weak (Supplementary Fig 3c), leaving exposed ssDNA susceptible to oxidative degradation^39^. Nonetheless, only in the presence of both a repA deletion and PIWI mutation, N-del/D663A/D738A, is DNA degradation completely lost. When a non-target plasmid with no complementarity to the supplied guides was incubated with the enzymes, fewer lower molecular weight products were generated by NgAgo relative to that when incubated with target plasmid containing a. While this result suggests off-target or guide-independent activity, this activity is reduced relative to guided cleavage as evidenced by fewer degradation products (Fig. 2f). That is, NgAgo-induced DNA degradation was also both target specific and non-specific, consistent with proposed pAgo models of non-specific DNA ‘chopping’ for guide acquisition and enhanced specific cleavage of complementary sequences^38^.

### NgAgo has specific *in vivo* activity at plasmid and genomic loci in bacteria

Next, we tested whether NgAgo can be programmed to target DNA *in vivo*. We chose *E.coli* instead of mammalian cells as our model because NgAgo, like most pAgos, lacks helicase activity needed to separate DNA strands for pAgo recognition and nicking of complementary sequences^18^,. The rapid rate of bacterial DNA replication increases the abundance of accessible unpaired DNA targets for NgAgo actitivy. Additionally, *E.coli* lack histones, which are known to inhibit pAgo activity^22^.

Studies have reproducibly demonstrated an ability of NgAgo to reduce gene expression^26,28^ and have suggested RNA cleavage as a possible mechanism. However, two alternative hypotheses could also explain this phenomena: (i) NgAgo cuts DNA leading to poor expression, and (ii) NgAgo inhibits transcription by tightly binding DNA. To distinguish between these three hypotheses, we created a two-plasmid system that harbors an inducible NgAgo expression cassette on one plasmid and another that serves as a target harboring a transcriptionally inactive pseudogene target, *mNeonGreen*, and a selectable marker or essential gene under selective conditions, *cat* (Fig. 3a). NgAgo was expressed in cells with both these plasmids and transformed with phosphorylated guide ssDNA (P-ssDNA) targeting different strands of *mNeonGreen*, including forward (FW, sense/coding), reverse (RV, antisense/non-coding), both FW and RV, or without a guide. After transformation, these cells were streaked on selective media (Fig. 3b). When guides were targeted to the transcriptionally silent *mNeonGreen* (Supplementary Fig. 5), fewer than half the colony forming units were observed relative to unguided controls (Fig. 3c). Control studies with either guides alone or NgAgo alone did not identify any cell toxicity, suggesting that the reduction in survival was due to NgAgo activity (Supplementary Figs.6 and 7). As similar results were obtained regardless of strand targeted and the target produced no RNA, NgAgo must interact at the DNA level. One possible mechanism is plasmid curing and loss of the selective marker through cleavage of the test plasmid, in agreement with our *in vitro* (Supplementary Figs 3 and 8) and cell-free studies (Fig 2). Using BFP in place of NgAgo does not reduce survival when incubated with guides complementary to the pseudogene *mNeonGreen* (Fig. 3c), confirming the survival reduction effect requires NgAgo expression. Finally, this effect is target specific. When targeted to an absent locus (*tetA*), there were no significant changes in the number of surviving colonies relative to unguided controls (Fig. 3c). This assay only quantifies activity relative to an unguided control and as such cannot measure off-target activity present in unguided controls. However, the reduction of survival in a guide- and target-dependent manner suggests that NgAgo has the capacity for targeted DNA endonuclease activity *in vivo* in *E. coli*.

**Figure 3.**
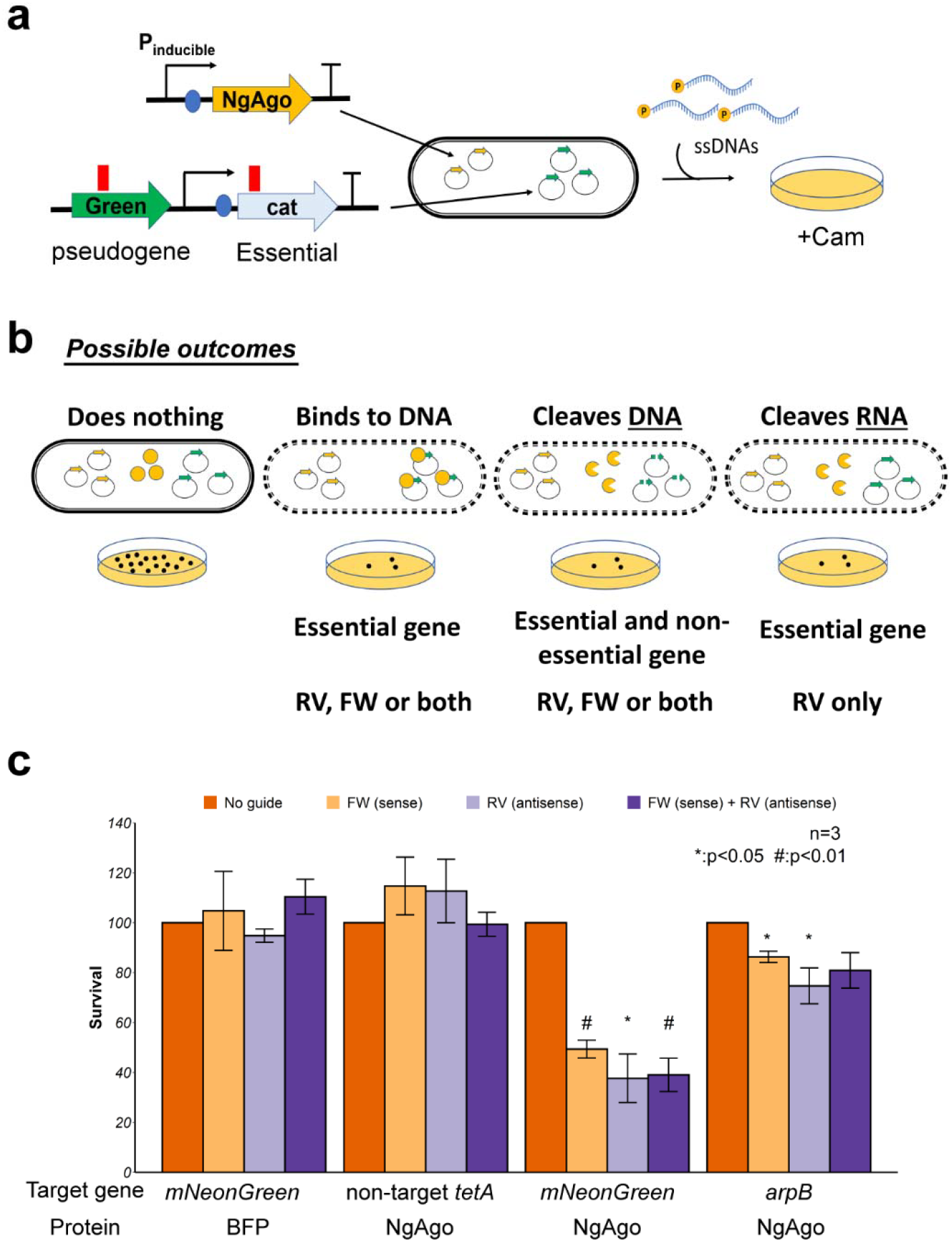
NgAgo can be programmed to target DNA in *E. coli*. **a**, Workflow of testing NgAgo function in *E. coli*. Two plasmids system used to test the function of NgAgo. One plasmid harbors NgAgo driven by T7 inducible promoter while the other low-copy plasmid serves as the target of NgAgo, including an untranscribed pseudogene, mNeonGreen. **b**. Four possible outcomes relative to an unguided control including no interaction, DNA binding, DNA cleaving, and RNA binding/cleaving, reveal the function of NgAgo. **c**, Survival rate targeting a pseudogene (mNeonGreen) on the plasmid or targeting a nonessential gene (arpB) in the genome with NgAgo or BFP control.

To confirm that the reduced survival is not limited to targets on the plasmid, we also targeted a genomic locus, *arpB*. *arpB* is a non-essential pseudogene that is interrupted by a stop codon^40^. Since *arpB* RNA is not required for survival (i.e., the arpB mutant is nonlethal), RNA cleavage would not reduce survival. However, double stranded DNA breaks in *E. coli* are lethal due to inhibited genome replication^41^. As targeting *arpB* did reduce survival (Fig. 3c), this suggests NgAgo also cleaves genomic DNA, consistent with our plasmid cleavage results.

Next, we asked if repA and PIWI domains are required for targeting in *E.coli* by evaluating the ability of different variants to target *mNeonGreen*. Our results showed that the PIWI mutant (D663A/D738A) and truncated repA deletion (N-del) lost the ability to reduce survival (Supplementary Fig. 9), suggesting the process of targeting and DNA cleavage was disrupted. Moreover, PIWI mutation enhanced survival activity via unknown mechanisms (Supplementary Fig. 9), potentially via its interactions with guide and other proteins^19^. Nonetheless, both intact repA and PIWI domains were required for targeted NgAgo activity.

### DNA-cleaving domains are needed for NgAgo programmable genome editing in bacteria

Since we have shown that NgAgo can cleave DNA *in vitro* and in *E.coli*, we asked whether this activity was essential for the reproducible gene editing by NgAgo observed in other prokaryotes^19^. To test for NgAgo gene editing activity, we created a kanamycin sensitive MG1655 (DE3) strain harboring a cassette composed of a *kanR* resistance gene lacking an RBS and promoter and a *mNeonGreen* gene flanked by two double terminators (Fig. 4a). This arrangement prevented any KanR/mNeonGreen expression from transcription read-through and translation from upstream and downstream genes. We then provided a donor plasmid with a truncated *mNeonGreen*, a constitutive promoter, an RBS and a truncated *kanR*, which is also KanR^−^ but can recombine with our locus to create a KanR^+^ phenotype (Fig. 4a). As DNA breaks in *E.coli* are lethal, repair via recombination should increase the number of KanR^+^ transformants if NgAgo induces DNA cleavage. We validated this system with CRISPR/Cas9, which showed a 4-fold enhancement in recombination efficiency (Supplementary Fig. 10).

**Figure 4.**
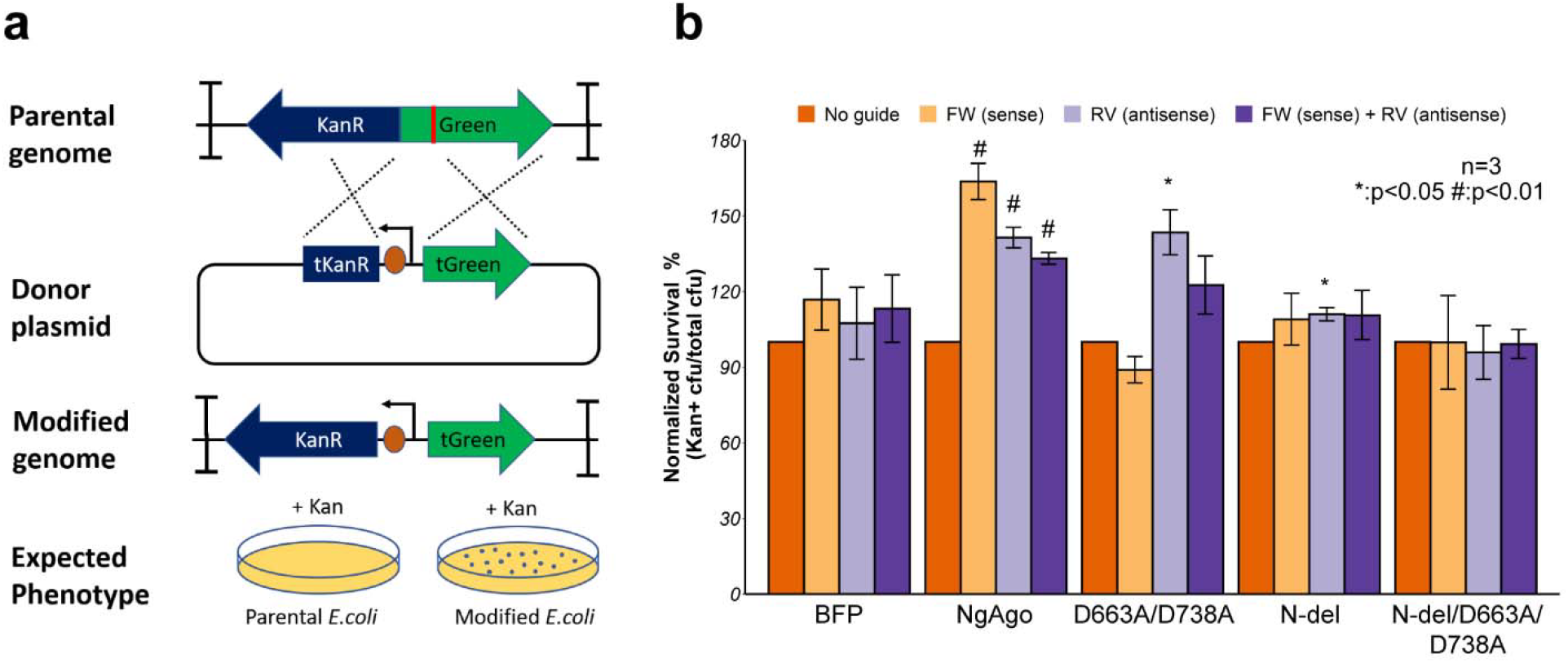
NgAgo enhances gene-editing via ◻-red-mediated homologous recombination in *E.coli*. **a,** Design of gene-editing assay in MG1655 (DE3). *KanR* and *mNeonGreen* (Green) cassette without promoter and RBS, flanked by two double terminators, is integrated in MG1655 (DE3). Donor plasmid with truncated *mNeonGreen* (tGreen) encodes a nonfunctional truncated *KanR* (tKanR). Guide was transformed to target the *mNeonGreen* (red line). After successful gene editing, modified genome has a functional KanR cassette, enabling survival in Kan selective plate. **b**, NgAgo variants enhance gene editing efficiency with ~1 microgram of guide(s) relative to an unguided control while blue fluorescent protein (BFP) control has no enhancement with guides. Error bars are the standard errors generated from three replicates. Statistically significant results are indicated with * (p-value< 0.05, paired t-test).

Wildtype NgAgo increased homologous recombination efficiency when provided with FW, RV, and both guides compared with an unguided control (Fig. 4b), demonstrating that guide-dependent NgAgo activity can enhance gene editing. In contrast, a BFP protein control showed no statistically significant enhancement in recombination compared to the unguided control (Fig. 4b). The PIWI mutant of NgAgo, D663A/D738A, displayed reduced but some statistically significant enhancement in homologous recombination; however, this was only true for one of the guides tested. The PIWI mutant displayed no significant enhancement of recombination with the FW or both guides (Fig. 4b). While the mechanism behind this pattern is unclear, these data suggest that the catalytic tetrad within the PIWI domain is not essential for enhanced homologous recombination under some conditions, in agreement with other published studies^19^. The N-del mutant of NgAgo lacking the repA domain displayed even weaker statistically significant enhancement in homologous recombination above unguided controls (11%) in the presence of the RV guide only (Fig. 4b). The N-del/D663A/D738A catalytic mutant showed no increase in gene editing activity in the presence of FW, RV, or both guides compared to an unguided control. This trend in homologous recombination enhancement is consistent with our observed DNA endonuclease activities (Fig 2e) suggesting that the DNA endonuclease activity mediated by the repA and PIWI domains is essential for enhanced homologous recombination and gene editing.

## DISCUSSION

NgAgo has been subject to intense debate in the literature in recent years^23,24,26,27,42^. Although previous studies suggested that refolded NgAgo does not cut DNA *in vitro*^18,31^, consistent with our findings, we establish that soluble NgAgo can, in fact, cleave DNA *in vitro*. That is, refolded NgAgo, which has been historically studied due to the poor soluble expression of this halophilic enzyme, may not be an accurate assessment of NgAgo activities. However, when soluble protein is concentrated and isolated, there is indeed some capacity for nonspecific or guide-independent DNA cleavage as we have demonstrated *in vitro*. Moreover, this behavior may be salt dependent, reflecting the halophilic lifestyle of the native host; NgAgo expressed from cells grown with LB Lennox showed no activity in our hands (data not shown) relative to that produced from cells grown on LB Miller (this work). Our parallel studies in cell-free expression systems that allow for control of salt conditions and lack potentially contaminating endonuclease expression confirm this observation. Most importantly, we generated a catalytically dead N-del/D663A/D738A mutant making it unlikely that the detected activity is the result of sample contamination.

NgAgo activity is mediated not only by the PIWI domain, like canonical pAgos, but also an uncharacterized and previously unrecognized accessory repA or single-stranded DNA binding domain fused to the N-terminus that appears common among halophilic pAgos (Fig 1c). Our work is the first report to suggest a role for this domain in NgAgo function and may be another source of the ongoing literature debate. Previously studied ‘catalytic’ mutants left this domain intact and were unable to detect a change in NgAgo function suggesting sample contamination or inactivity^31^. However, this and growing evidence from the literature^18–20^ suggest that accessory proteins and domains may be essential for pAgo function. As homologous accessory proteins from heterologous hosts can mediate function^18,19^, we investigated whether *in vivo* cleavage, as observed via cell survival and DNA recombination efficiency, would be induced by NgAgo and its mutants. Not only were these assay results consistent with DNA cleavage, but they also importantly suggested an ability to target specific gene loci via single-stranded 5’P DNA guides. Our work here underscores the role of unrecognized accessory proteins, supplied via the expression host, and a need to characterize these proteins to more accurately assess pAgo activity.

Finally, our results provide supporting evidence to encourage the development of NgAgo for gene-editing. When provided with homologous target and donor sequences, NgAgo can enhance homologous recombination. Much like other pAgos, the PIWI domain participates in DNA editing in prokaryotes as shown here and by Fu *et al*^19^. Moreover, without repA, PIWI mutants of NgAgo exhibit reduced cleavage activity with a concomitant reduction in homologous recombination efficiency. Both the repA deletion and the PIWI mutation (N-del/D663A/D738) are needed to fully abolish catalytic and gene-editing functions. In the presence of both functional domains, NgAgo can effectively enhance homologous recombination by inducing a double stranded break at a targeted region. Despite the programmable DNA-cleaving ability of NgAgo, there remain several challenges to its development as a robust tool for gene-editing applications: guide-independent or off-target cleavage, unknown accessory proteins needed for function, poor expression, salt dependence, and potentially low activity in eukaryotic hosts. Nonetheless, further insight may lead to protein engineering strategies to overcome these hurdles and develop NgAgo as a robust tool for gene-editing.

## Conclusion

Based on the above findings, we conclude that NgAgo is a novel DNA endonuclease that belongs to an unrecognized class of pAgos defined by a characteristic repA domain. NgAgo uses both a well-conserved catalytic tetrad in PIWI and a novel uncharacterised repA domain to cleave DNA. This cleavage activity is essential to enhancing gene-editing efficiency in prokaryotes. Despite the challenges of NgAgo, our work establishes innovative approaches to probe NgAgo activity (and that of other pAgos) and identifies critical protein features for its development as a next generation synthetic biology tool.

## Acknowledgment

We are grateful to Dr. Xin Ge (University of California, Riverside) and Dr. Kristala J. Prather (Massachusetts Institute of Technology) for providing pET32a-GST-ELP64 plasmid and MG1655 (DE3), respectively. We also thank Dr. Mathew Tantama (Purdue University) for providing pBAD-mTagBFP2 plasmid.

## Author contributions

K.V.S., F.G., and K.Z.L. designed the experiments. K.Z.L., M.A.M., A.K., A.L., K. F., and P.P. conducted and analyzed the experiments. K.V.S., F.G., and K.Z.L. supervised research and experimental design. K.V.S, K.Z.L, M.A.M, and F.G. wrote the manuscript.

## MATERIAL AND METHODS

### Strains and plasmids

*E. coli* strains and plasmids used in this study are listed in Table 1. Cloning was carried out according to standard practices^43^ with primers, template, and purpose listed in Supplementary Table 5. Plasmids were maintained in *E. coli* DH5α. NgAgo variants (wildtype, D663A/D738A, N-del, and repA with GST or His tag) that were used for *in vitro* activity assays were cloned into an IPTG-inducible T7 plasmid, pET32a-GST-ELP64. MG1655 (DE3) *atpI*::KanR-mNeonGreen was generated using recombineering^44^ via donor plasmid pTKDP-KanR-mNeonGreen-hph. For gene-editing/recombination studies^45^, p15-KanR-PtetRed was used as a donor plasmid (Table 1).

**Table 1.**
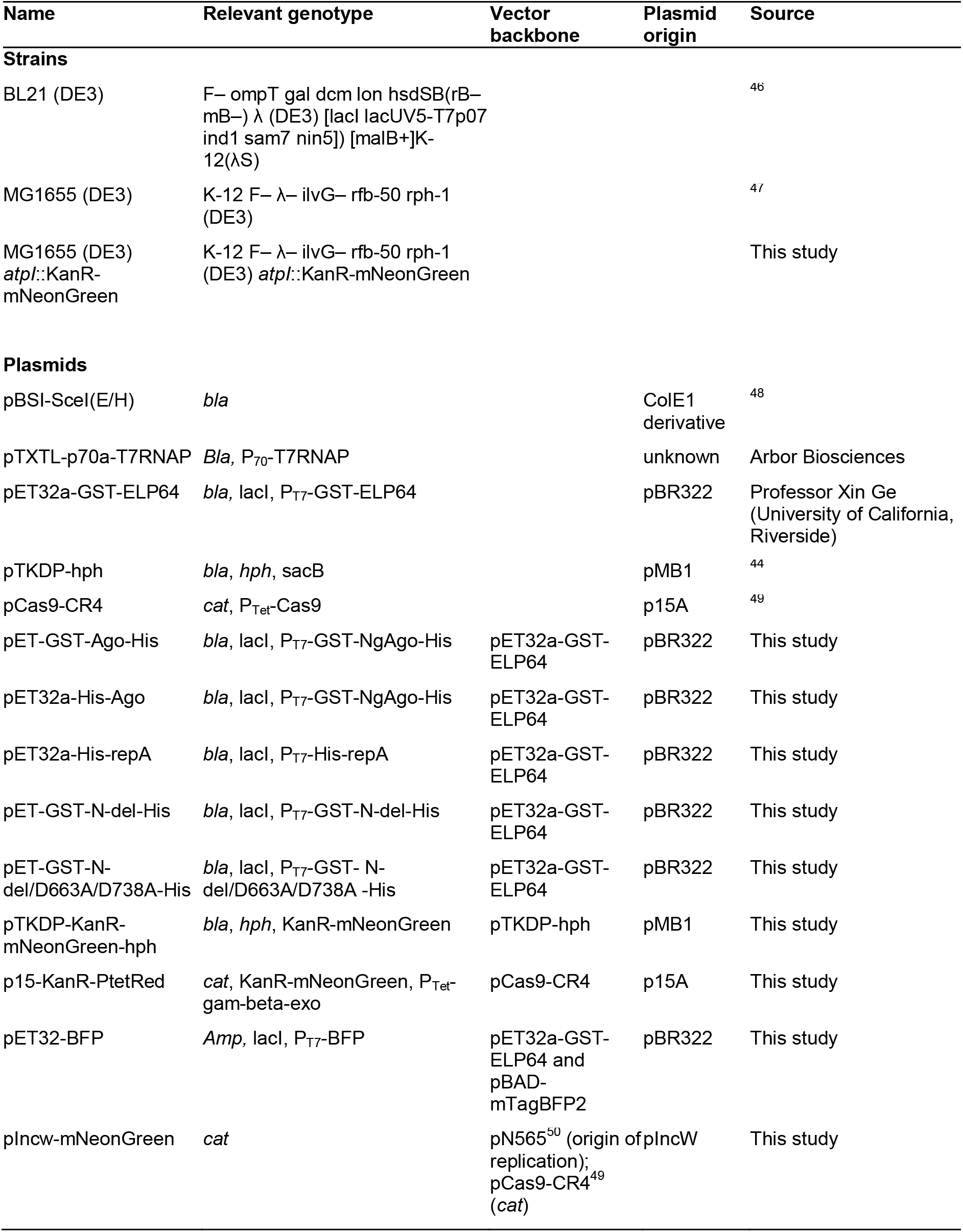
Strains and Plasmids.

### NgAgo expression and purification

GST-NgAgo or His-NgAgo variants were expressed in BL21 (DE3) with 100 μg/ml ampicillin. 5 mL cultures started from single colonies were grown for 16 hours before subculturing in 100 ml of LB Miller containing ampicillin. Expression was induced with 0.1 mM IPTG at OD_600_ = 0.5 for either 4 hours at 37 °C or overnight at 22 °C overnight before harvesting the cells at 7500 rpm (11,500 g) at 4 °C for 5 minutes. The cell pellet was resuspended in TN buffer (10 mM Tris and 100mM NaCl, pH 7.5) and lysed via sonication at a medium power setting (~50 W) in 10 s intervals, with intervening 10 s incubations on ice to reduce heat denaturation. Cell lysates were then clarified at 12000 rpm at 4 °C for 30 minutes. The supernatant was collected as a soluble protein fraction. Both soluble and insoluble (cell pellet) fractions were purified via His-IDA nickel column (Clontech Laboratories, Mountain View, CA. Cat. No: 635657) according to the manufacturer instructions. Insoluble NgAgo protein was refolded on the column after denaturation with guanidium chloride according to manufacturer instructions. GST-tagged NgAgo variants were purified by glutathione agarose (Thermo Fisher Scientific, Waltham, MA. Cat. No: 16100) according to the manufacturer protocol.

### Cell-free expression of NgAgo and activity assay

Cell-free TXTL reactions contained 5’ phosphorylated DNA guides, Chi6 oligos, IPTG, plasmids encoding T7RNA polymerase (pTXTL-p70a-T7RNAP) and NgAgo variants, including wildtype, D663A/D738A, repA, N-del, and N-del/D663A/D738A (Table 2). Reactions were incubated at 29 °C for 20 hours to promote NgAgo expression before being supplemented to 125 mM NaCl and incubating at 37 °C for folding for 24 hours. MgCl_2_ to a final concentration of 62.5 μM was then added along with target or non-target plasmid for reaction at 37 °C for an hour. RNase A (70 ng or >490 units) (Millipore Sigma, Burlington, MA. Cat. No: R6513-10MG) was then added to each reaction to remove transcribed RNA at 37 °C for 10 minutes. The reaction mixtures were then mixed with 0.5% SDS to dissociate any proteins and 6X loading dye before gel electrophoresis. The gel was visualized under a blue light (Azure Biosystems, Dublin, CA. Azure c400).

**Table 2.**
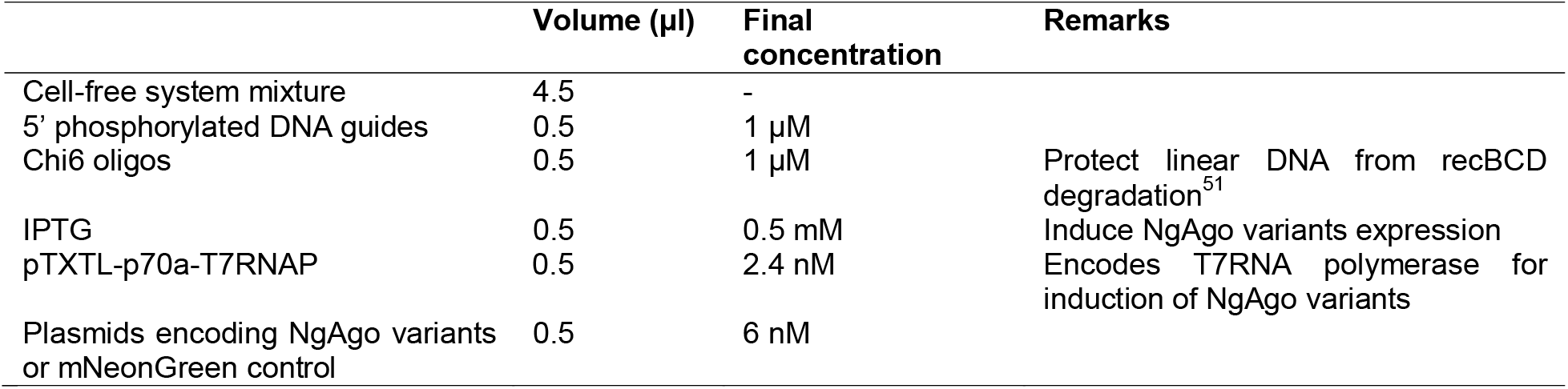
Materials for NgAgo variants production by cell-free system.

### Survival assay

BL21 (DE3) was transformed with target plasmid pIncw-mNeonGreen and NgAgo expression plasmid and made electrocompetent. Electrocompetent cells were transformed with either no guides or 1 μg total of FW, RV, both guides and plated on ampicillin and chloramphenicol selective LB Miller agar plate with 0.1 mM IPTG before 16-20 hours incubation at 37 °C. Colonies were counted to measure survival rate of transformants. The unguided control was normalized to 100% and guided-treatments were normalized to the unguided control.

### Gene-editing assay

MG1655 (DE3) *atpI*::KanR-mNeonGreen was transformed with pET-GST-NgAgo-His (to induce DNA cleavage) and p15-KanR-PtetRed (for lambda-red recombinase expression and to provide donor DNA for repair) and made electrocompetent. Electrocompetent cells were transformed with either no guides or one 1.2 μl of 100 μM total of FW, RV, both guides and incubated in LB Miller with ampicillin, chloramphenicol, and IPTG for an hour. These cultures were then diluted ten-fold in LB Miller containing ampicillin (working concentration: 100 μg/ml), chloramphenicol (working concentration: 25 μg/ml), IPTG (working concentration: 0.1mM), and anhydrotetracycline (aTc) (working concentration: 50 μg/ml), incubated until OD_600_ = 0.2 before plating with and without kanamycin (working concentration: 50 μg/ml). Colony forming units (CFU) were counted after 16-20 hours incubation at 37 °C. The unguided control was normalized to 100% and guided-treatments were normalized to the unguided control.

### Phyre 2 and HHpred analysis

NgAgo protein (IMG/M Gene ID: 2510572918) was analyzed via Phyre 2^32^ with normal mode on 2018 November 19. The normal mode pipeline involves detecting sequence homologues, predicting secondary structure and disorder, constructing a hidden Markov model (HMM), scanning produced HMM against library of HMMs of proteins with experimentally solved structures, constructing 3D models of NgAgo, modelling insertions/deletions, modelling of amino acid sidechains, submission of the top model, and transmembrane helix and topology prediction^32^. NgAgo was analyzed via HHpred^33,34^ (https://toolkit.tuebingen.mpg.de/#/tools/hhpred) on 2018 November 27. The parameters for HHpred are HHblits=>uniclust30_2018_08 for multiple sequence alignment (MSA) generation method, 3 for maximal number of MSA generation steps, 1e-3 for E-value incl. threshold for MSA generation, 0% for minimum sequence identity of MSA hits with query, 20% for minimum coverage of MSA hits, during_alignment for secondary structure scoring, local for alignment mode, off for realign with MAC, 0.3 for MAC realignment threshold, 250 for number of target sequences, and 20% for minimum probability in hit list.

### Phylogenetic analysis

BLAST was used to compare NgAgo protein sequence with all the isolates in the database via the IMG/M server (https://img.jgi.doe.gov/). Representative full-length Argonautes with a repA domain were used to represent each species. Selected pAgos with repA domains and some well-characterized pAgos were compared, and the midpoint rooted tree was generated via the server http://www.genome.jp/tools-bin/ete with unaligned input type, mafft_default aligner, no alignment cleaner, no model tester, and fasttree_default Tree builder parameters. The nwk output file was then used for phylogenetic tree generation in R with ggtree package.

## References

1. Hegge, J. W., Swarts, D. C. & van der Oost, J. Prokaryotic Argonaute proteins: novel genome-editing tools? Nature Reviews Microbiology 16, 5 (2018).

2. Swarts, D. C. et al. DNA-guided DNA interference by a prokaryotic Argonaute. Nature 507, 258–261 (2014).

3. Willkomm, S. et al. Structural and mechanistic insights into an archaeal DNA-guided Argonaute protein. Nature Microbiology 2, 17035 (2017).

4. Enghiad, B. & Zhao, H. Programmable DNA-guided artificial restriction enzymes. ACS Synthetic Biology 6, 752–757 (2017).

5. Kaya, E. et al. A bacterial Argonaute with noncanonical guide RNA specificity. Proceedings of the National Academy of Sciences 113, 4057–4062 (2016).

6. Hauptmann, J. et al. Turning catalytically inactive human Argonaute proteins into active slicer enzymes. Nature Structural and Molecular Biology 20, 814 (2013).

7. Faehnle, C. R., Elkayam, E., Haase, A. D., Hannon, G. J. & Joshua-Tor, L. The making of a slicer: activation of human Argonaute-1. Cell Reports 3, 1901–1909 (2013).

8. Kwak, P. B. & Tomari, Y. The N domain of Argonaute drives duplex unwinding during RISC assembly. Nature Structural & Molecular Biology 19, 145 (2012).

9. Ma, J.-B. et al. Structural basis for 5′-end-specific recognition of guide RNA by the A. fulgidus Piwi protein. Nature 434, 666 (2005).

10. Künne, T., Swarts, D. C. & Brouns, S. J. J. Planting the seed: target recognition of short guide RNAs. Trends in Microbiology 22, 74–83 (2014).

11. Lingel, A., Simon, B., Izaurralde, E. & Sattler, M. Nucleic acid 3′-end recognition by the Argonaute2 PAZ domain. Nature Structural and Molecular Biology 11, 576 (2004).

12. Ma, J.-B., Ye, K. & Patel, D. J. Structural basis for overhang-specific small interfering RNA recognition by the PAZ domain. Nature 429, 318 (2004).

13. Sheng, G. et al. Structure-based cleavage mechanism of Thermus thermophilus Argonaute DNA guide strand-mediated DNA target cleavage. Proceedings of the National Academy of Sciences 111, 652–657 (2014).

14. Wang, Y. et al. Structure of an argonaute silencing complex with a seed-containing guide DNA and target RNA duplex. Nature 456, 921 (2008).

15. Hur, J. K., Zinchenko, M. K., Djuranovic, S. & Green, R. Regulation of Argonaute slicer activity by guide RNA 3’end interactions with the N-terminal lobe. Journal of Biological Chemistry jbc–M112 (2013).

16. Swarts, D. C. et al. The evolutionary journey of Argonaute proteins. Nature Structural & Molecular Biology 21, 743–753 (2014).

17. Ryazansky, S., Kulbachinskiy, A. & Aravin, A. A. The Expanded Universe of Prokaryotic Argonaute Proteins. mBio 9, (2018).

18. Hunt, E. A., Evans Jr, T. C. & Tanner, N. A. Single-stranded binding proteins and helicase enhance the activity of prokaryotic argonautes in vitro. PloS One 13, e0203073 (2018).

19. Fu, L. et al. The prokaryotic Argonaute proteins enhance homology sequence-directed recombination in bacteria. Nucleic Acids Research 47, 3568–3579 (2019).

20. Jolly, S. M. et al. Thermus thermophilus Argonaute Functions in the Completion of DNA Replication. Cell 182, 1545–1559.e18 (2020).

21. Swarts, D. C. et al. Argonaute of the archaeon Pyrococcus furiosus is a DNA-guided nuclease that targets cognate DNA. Nucleic Acids Research 43, 5120–5129 (2015).

22. Zander, A. et al. Guide-independent DNA cleavage by archaeal Argonaute from Methanocaldococcus jannaschii. Nature Microbiology 2, 17034 (2017).

23. Cyranoski, D. Authors retract controversial NgAgo gene-editing study. Nature News (2017) doi:10.1038/nature.2017.22412.

24. Javidi-Parsijani, P. et al. No evidence of genome editing activity from Natronobacterium gregoryi Argonaute (NgAgo) in human cells. PLoS One 12, 14 (2017).

25. Wu, Z. et al. NgAgo-gDNA system efficiently suppresses hepatitis B virus replication through accelerating decay of pregenomic RNA. Antiviral Research 145, 20–23 (2017).

26. Burgess, S. et al. Questions about NgAgo. Protein & Cell 7, 913–915 (2016).

27. Khin, N. C., Lowe, J. L., Jensen, L. M. & Burgio, G. No evidence for genome editing in mouse zygotes and HEK293T human cell line using the DNA-guided Natronobacterium gregoryi Argonaute (NgAgo). PloS One 12, e0178768 (2017).

28. Qin, Y. Y., Wang, Y. M. & Liu, D. NgAgo-based fabp11a gene knockdown causes eye developmental defects in zebrafish. Cell Research 26, 1349–1352 (2016).

29. Elcock, A. H. & McCammon, J. A. Electrostatic contributions to the stability of halophilic proteins. Journal of Molecular Biology 280, 731–748 (1998).

30. Tadeo, X. et al. Structural basis for the amino acid composition of proteins from halophilic archea. PLoS Biology 7, e1000257 (2009).

31. Sunghyeok, Y. et al. DNA-dependent RNA cleavage by the Natronobacterium gregoryi Argonaute. BioRxiv 101923 (2017).

32. Kelley, L. A., Mezulis, S., Yates, C. M., Wass, M. N. & Sternberg, M. J. E. The Phyre2 web portal for protein modeling, prediction and analysis. Nature Protocols 10, 845–858 (2015).

33. Zimmermann, L. et al. A completely Reimplemented MPI bioinformatics toolkit with a new HHpred server at its Core. Journal of Molecular Biology 430, 2237–2243 (2018).

34. Söding, J., Biegert, A. & Lupas, A. N. The HHpred interactive server for protein homology detection and structure prediction. Nucleic Acids Research 33, W244–W248 (2005).

35. Flynn, R. L. & Zou, L. Oligonucleotide/oligosaccharide-binding fold proteins: a growing family of genome guardians. Critical Reviews in Biochemistry and Molecular Biology 45, 266–275 (2010).

36. Müller-Santos, M. et al. First evidence for the salt-dependent folding and activity of an esterase from the halophilic archaea Haloarcula marismortui. Biochimica et Biophysica Acta (BBA)-Molecular and Cell Biology of Lipids 1791, 719–729 (2009).

37. Marshall, R. et al. Rapid and Scalable Characterization of CRISPR Technologies Using an E. coli Cell-Free Transcription-Translation System. Molecular Cell 69, 146–157.e3 (2018).

38. Swarts, D. C. et al. Autonomous Generation and Loading of DNA Guides by Bacterial Argonaute. Molecular Cell 65, 985–998 (2017).

39. Bell, J. C., Liu, B. & Kowalczykowski, S. C. Imaging and energetics of single SSB-ssDNA molecules reveal intramolecular condensation and insight into RecOR function. eLife 4, e08646 (2015).

40. Goodall, E. C. A. et al. The Essential Genome of Escherichia coli K-12. mBio 9, (2018).

41. Simmons, L. A. et al. Comparison of Responses to Double-Strand Breaks between Escherichia coli and Bacillus subtilis Reveals Different Requirements for SOS Induction. Journal of Bacteriology 191, 1152–1161 (2009).

42. Wu, Z. et al. NgAgo-gDNA system efficiently suppresses hepatitis B virus replication through accelerating decay of pregenomic RNA. Antiviral Research 145, 20–23 (2017).

43. Sambrook, J., Fritsch, E. F. & Maniatis, T. Molecular cloning: a laboratory manual. (Cold Spring Harbor Laboratory Press, 1989).

44. Tas, H., Nguyen, C. T., Patel, R., Kim, N. H. & Kuhlman, T. E. An integrated system for precise genome modification in Escherichia coli. PloS One 10, e0136963 (2015).

45. Jiang, Y. et al. Multigene editing in the Escherichia coli genome via the CRISPR-Cas9 system. Applied and Environmental Microbiology 81, 2506–2514 (2015).

46. Wood, W. B. Host specificity of DNA produced by Escherichia coli: bacterial mutations affecting the restriction and modification of DNA. Journal of Molecular Biology 16, 118–IN3 (1966).

47. Tseng, H.-C., Martin, C. H., Nielsen, D. R. & Prather, K. L. J. Metabolic engineering of Escherichia coli for enhanced production of (R)-and (S)-3-hydroxybutyrate. Applied and Environmental Microbiology 75, 3137–3145 (2009).

48. Niu, Y., Tenney, K., Li, H. & Gimble, F. S. Engineering variants of the I-SceI homing endonuclease with strand-specific and site-specific DNA-nicking activity. Journal of Molecular Biology 382, 188–202 (2008).

49. Reisch, C. R. & Prather, K. L. J. The no-SCAR (Scarless Cas9 Assisted Recombineering) system for genome editing in Escherichia coli. Scientific Reports 5, 15096 (2015).

50. Rhodius, V. A. et al. Design of orthogonal genetic switches based on a crosstalk map of σs, anti-σs, and promoters. Molecular Systems Biology 9, 702 (2013).

51. Marshall, R., Maxwell, C. S., Collins, S. P., Beisel, C. L. & Noireaux, V. Short DNA containing χ sites enhances DNA stability and gene expression in E. coli cell-free transcription-translation systems. Biotechnology and Bioengineering 114, 2137–2141 (2017).

