## Supplemental Materials for "NgAgo DNA endonuclease activity enhances homologous recombination in *E. coli*"

**SUPPLEMENTAL NOTES**

***NgAgo has some nonspecific DNA cleavage activity***

We hypothesized that NgAgo generates random guides in the host via DNA chopping^1^, which co-purifies with NgAgo leading to apparent guide-independent activity *in vitro*. While we were able to confirm the presence of these random copurified guides (Supplementary Fig. 3c), we were unable to displace them with incubation at high temperature (55 °C) and reload with our target guides (reloading protocol). Subsequent testing had similar guide-independent cleavage activity with no evidence of increased linearized plasmid (Supplementary Fig. 3d). As refolded NgAgo had no cleavage activity, we used soluble NgAgo to study its function *in vitro* unless otherwise stated.

Previous studies have demonstrated that TtAgo can obtain random guides from the expression plasmid DNA via DNA chopping^1^. Thus, the observed guide-independent cleavage may indeed be guide-dependent as a result of chopping and subsequent guide loading with homologous DNA, which cannot be easily displaced as demonstrated in Supplementary Fig 3d. To examine this hypothesis, we completed the *in vitro* cleavage assay with a ‘related’ plasmid, pNCS-mNeonGreen (Supplementary Fig. 8a), and an ‘unrelated’ plasmid, p15-KanR (Supplementary Fig.8c). The unrelated plasmid, p15-KanR, shares no DNA homology with the NgAgo expression plasmid while the related plasmid, pNCS-mNeonGreen, has the same ampicillin resistance gene. NgAgo cleaved both related and unrelated plasmids independent of guide (Supplementary Fig. 8b and 8d), suggesting that the guide-independent cleavage activity of our purified NgAgo does not rely on pre-loaded DNA. These results confirmed that NgAgo has guide-independent cleavage activity *in vitro*, sharing similar properties with bacterial TtAgo^1^ and archaeal MjAgo^2^.

***Persistence of guide for* in vivo assays**

As our guide sequences are 5’P ssDNA, they are exogenously supplied via transformation. Guides are supplied before NgAgo induction for all *in vivo* assays in order to minimize DNA chopping due to guide-free NgAgo. However, guide molecules cannot replicate and are depleted through cell division and degradation. To assay their persistence and determine whether or not they persist long enough to encounter induced NgAgo, we transformed D4PA-labelled red ssDNA into cells expressing BFP. After 4 h when sufficient BFP has accumulated to be visible, labelled ssDNA still persists and is theoretically available for NgAgo binding (Supplementary Fig. 11).

***Validation of cell-free protein expression***

We used mNeonGreen as a reporter to indicate the successful protein production by the cell-free system (Supplementary Fig. 12).

***NgAgo may represent a new class of mesophilic pAgos***

To our knowledge, NgAgo is the first studied pAgo with a fused repA domain. This domain is essential for cleavage as evidenced in our cell-free *in vitro* studies and recombination assays. Interestingly, all repA domain-containing pAgos are from halophilic Archaea mesophiles, suggesting that the repA domain may have fused to pAgos in the last common ancestor of all halophiles. Moreover, single-stranded binding (SSB) proteins have been demonstrated to participate in and enhance pAgo activity^3^ suggesting that the repA domain at the N-terminus of NgAgo may be involved in the cleaving process without recruitment of endogenous SSB proteins. Further research, however, is needed to clarify the function of this repA domain.

**SUPPLEMENTAL METHODS**

***In vitro* activity assay**

For the reloading protocol, 5 µg purified NgAgo was mixed with 1 µg total of phosphorylated single-stranded DNA (P-ssDNA) targeting mNeonGreen (Supplementary Table 6) and incubated at 55 °C for an hour. 200-300 ng of substrate plasmid DNA (pNCS-mNeonGreen) was then added to the sample. The final volume of the reaction was 50 µl (working concentration: 20 mM Tris-Cl, 300 mM KCl, 500 μM MgCl_2_, and 2 mM DTT). The sample was then incubated at 37 °C for three hours. 0.8 units of Proteinase K (NEB, Ipswich, MA. Cat. No: P8107S) were added to the sample to digest the protein for 5 minutes at 37 °C. The nucleic acids were then isolated with the DNA Clean & Concentrator™-5 kit (Zymo Research, Irvine, CA. Cat. No: D4003T) according to manufacturer instructions and mixed with 6X loading dye containing SDS (Thermo Fisher S, Waltham, MA. Cat. No: R1151) before gel electrophoresis. The gel containing Sybrsafe (Thermo Fisher S, Waltham, MA. Cat. No: S33102) was visualized under a blue light (Azure Biosystems, Dublin, CA. Azure c400).

For our standard protocol, we incubated the same amount of guides and proteins at 37 °C for 30 minutes, and added the same amount of plasmid DNA (p15-KanR or pBSI-SceI(E/H)^4^) with 50 ul final volume (working concentration: 20mM Tris-Cl, 300mM NaCl, 250 uM MgCl_2_, and 2mM DTT). The samples were incubated at 37 °C for an hour before Proteinase K treatment. The rest of the procedure is the same as the reloading protocol.

**Electrophoretic mobility shift assay (EMSA)**

5 µg of purified N-del and repA were incubated with 1 µg of mNeonGreen ssDNA guide in 50ul in buffer (working concentration: 20 mM Tris-Cl, 300 mM KCl, 500 μM MgCl_2_, and 2 mM DTT) at 37 °C for an hour and treated with 0.8 units proteinase K for 5 minutes if needed before running with 20% TBE gel with 0.5X TBE buffer. Gels were stained with Sybr Gold (Thermo Fisher Scientific, Waltham, MA. Cat. No: S11494) before visualizing under a green fluorescent channel (Azure Biosystems, Dublin, CA. Azure c400). Positional marker 10/60 ladder (Coralville, IA. Cat. No: 51-05-15-01) was used in the EMSA assay.

**SUPPLEMENTARY DATA**

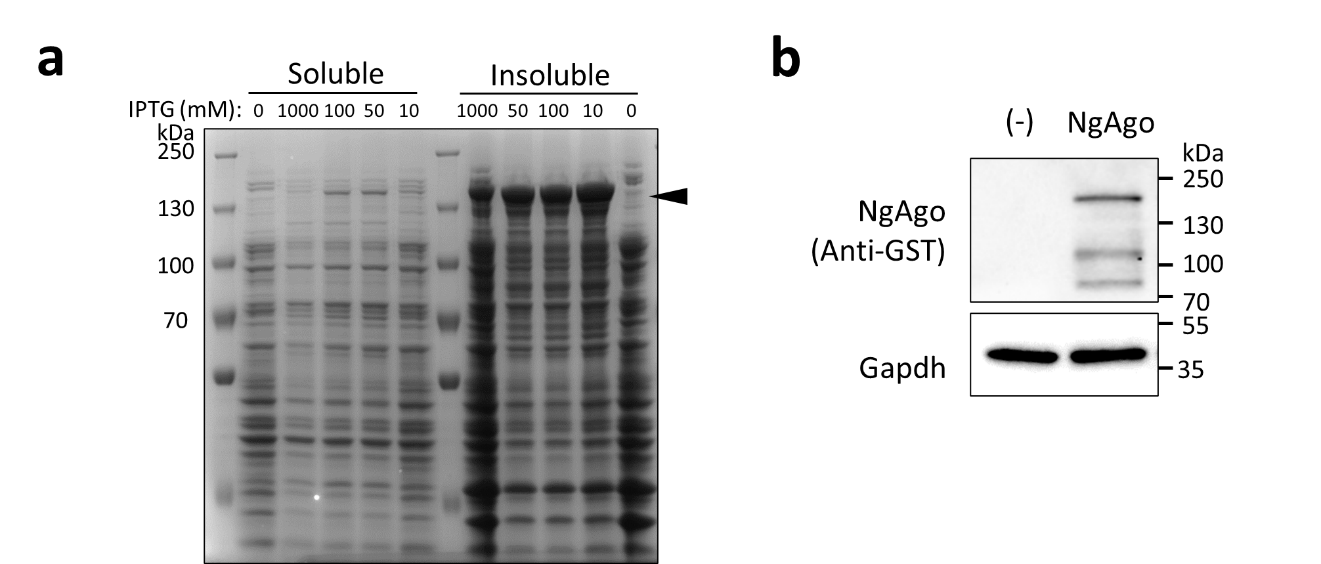

**Supplementary Figure 1 | Optimization of soluble NgAgo protein expression. a**, Different IPTG concentrations (1000 mM, 100 mM, 50 mM, and 10 mM) were used to induce GST-NgAgo expression in BL21 (DE3). Soluble and insoluble protein fractions were analyzed by SDS-PAGE to determine the optimal conditions for soluble NgAgo expression. **b**, Soluble GST-NgAgo expression with 100mM IPTG was probed with anti-GST antibody and a Gapdh internal control.

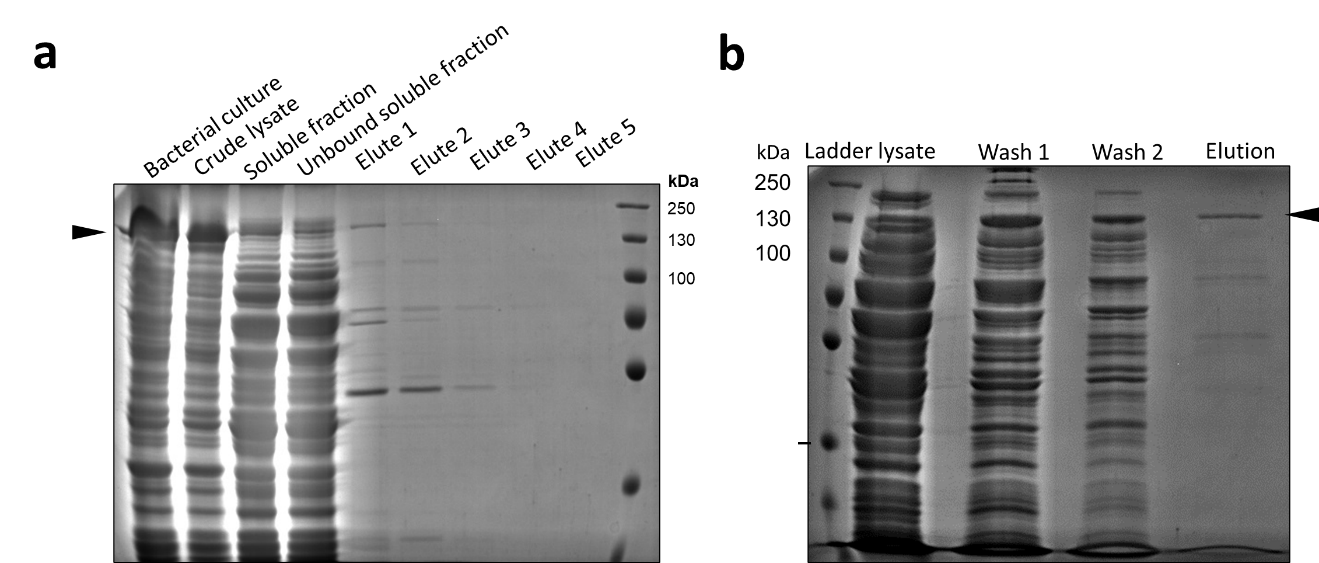

**Supplementary Figure 2 | SDS-PAGE analysis of His-tag purified NgAgo variants.** a, SDS-PAGE analysis of purified WT NgAgo from soluble fraction (sNgAgo). Elute 1 was used for *in vitro* assay. b, SDS-PAGE analysis of purified WT NgAgo from insoluble fraction after refolding (rNgAgo). Elution fraction was used for *in vitro* assay.

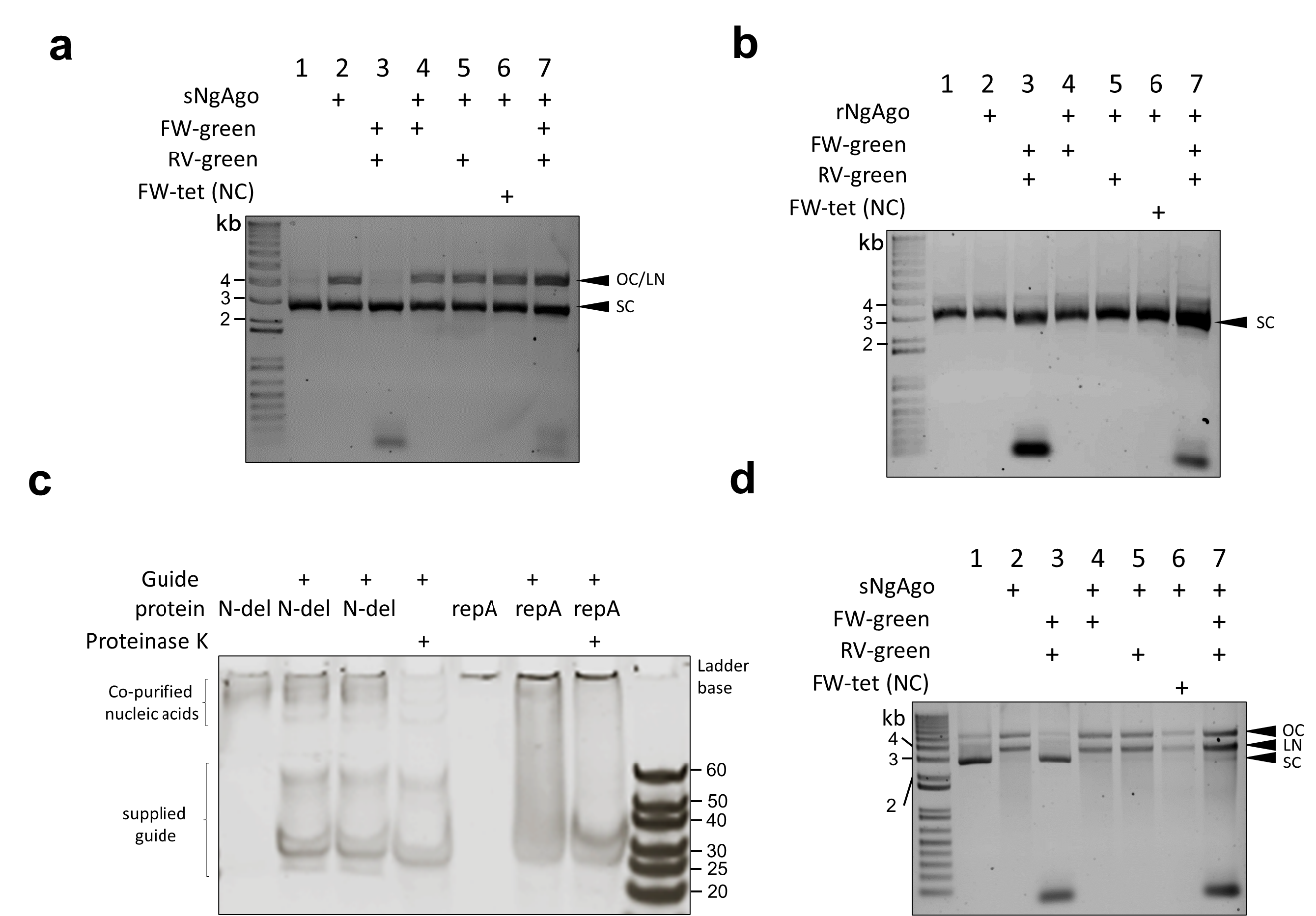

**Supplementary Figure 3 | Soluble NgAgo variants nick and cut plasmids DNA in vitro. a**, Soluble NgAgo (sNgAgo) nicks an cuts plasmids DNA regardless the presence of guide DNA. **b**, Refolded NgAgo, rNgAgo, has no effect on plasmids DNA regardless the presence of guide DNA. **c**, Electrophoretic mobility shift assay (EMSA) of N-del and repA domain with guides. N-del does not show band shifting while repA treatment shifts the bands. **d**, Soluble NgAgo (sNgAgo) nicks and cuts the plasmids DNA regardless the presence of guide DNA with Han’s guide-reloading protocol. OC: open circular; LN: linear; SC: supercoiled.

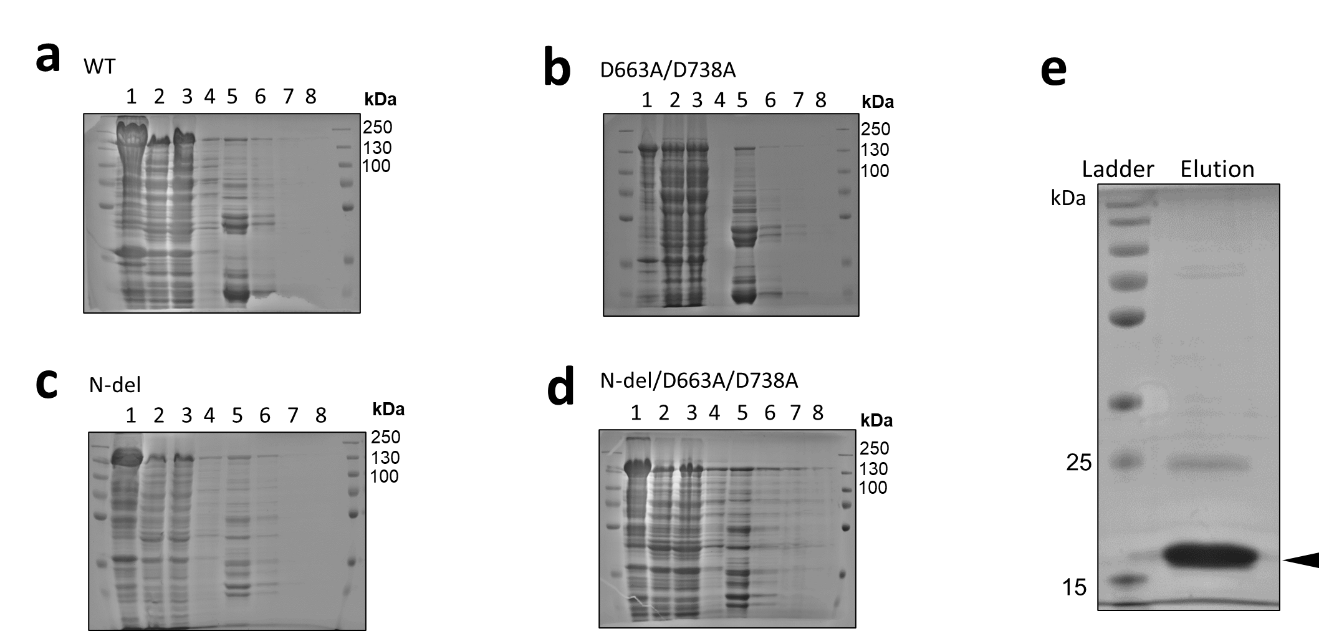

**Supplementary Figure 4 | SDS-PAGE analysis of GST-tag purified soluble NgAgo variants. a**, SDS-PAGE analysis of GST-tag purified WT NgAgo. **b**, SDS-PAGE analysis of GST-tag purified D663A/D738A. **c**, SDS-PAGE analysis of GST-tag purified N-del. **d**, SDS-PAGE analysis of GST-tag purified N-del/D663A/D738A. 1: whole cell lysate; 2: soluble fraction; 3: unbound soluble fraction; 4 washed fraction; 5-8: eluted fraction 1-4. **e**. His-tagged soluble repA

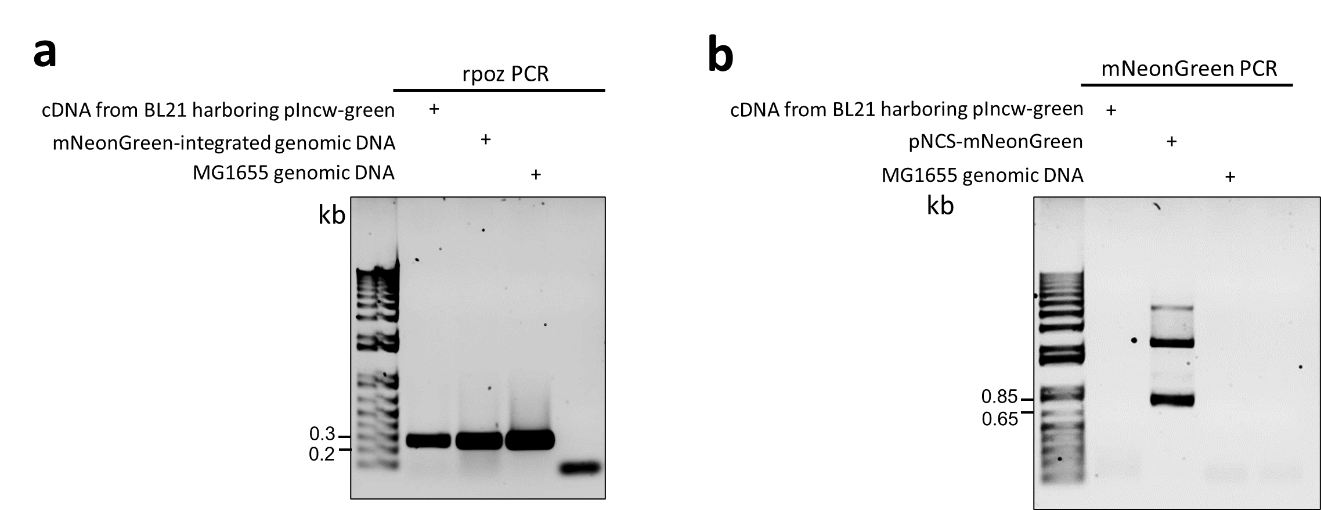

**Supplementary Figure 5 | mNeonGreen of pIncw-green is transcriptionally silent. a**, RNA polymerase subunit, *rpoz*, was amplified with cDNA from BL21 harboring pIncw-mNeonGreen, mNeonGreen-integrated genomic DNA, and WT genomic DNA. **b**, mNeonGreen was amplified with cDNA from BL21 harboring pIncw-mNeonGreen, pNCS-mNeonGreen plasmid DNA, and WT genomic DNA.

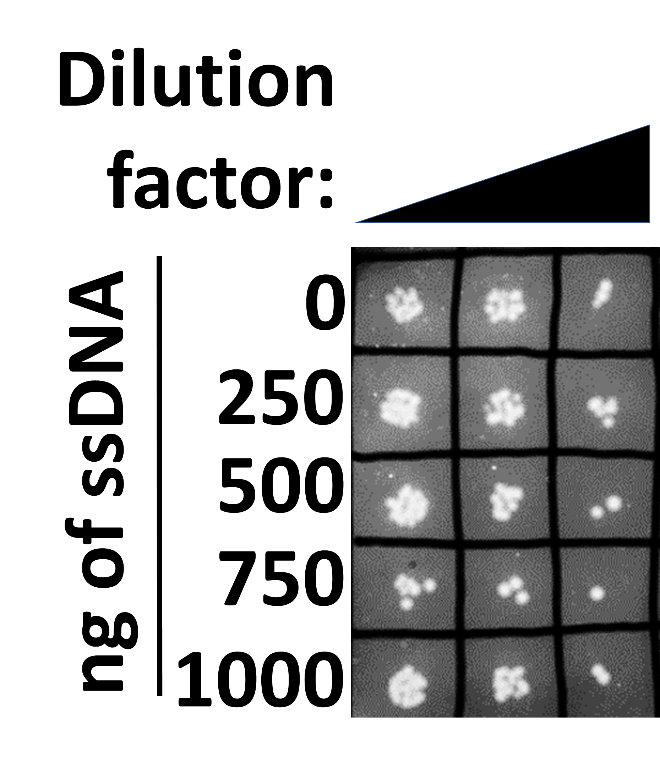

**Supplementary Figure 6 | ssDNA guides are non-toxic**. Different amounts of ssDNAs were transformed into BL21 and spot plated at varying dilutions (1000x, 2000x and 5000x). Ten microliter of each diluted sample was plated on LB.

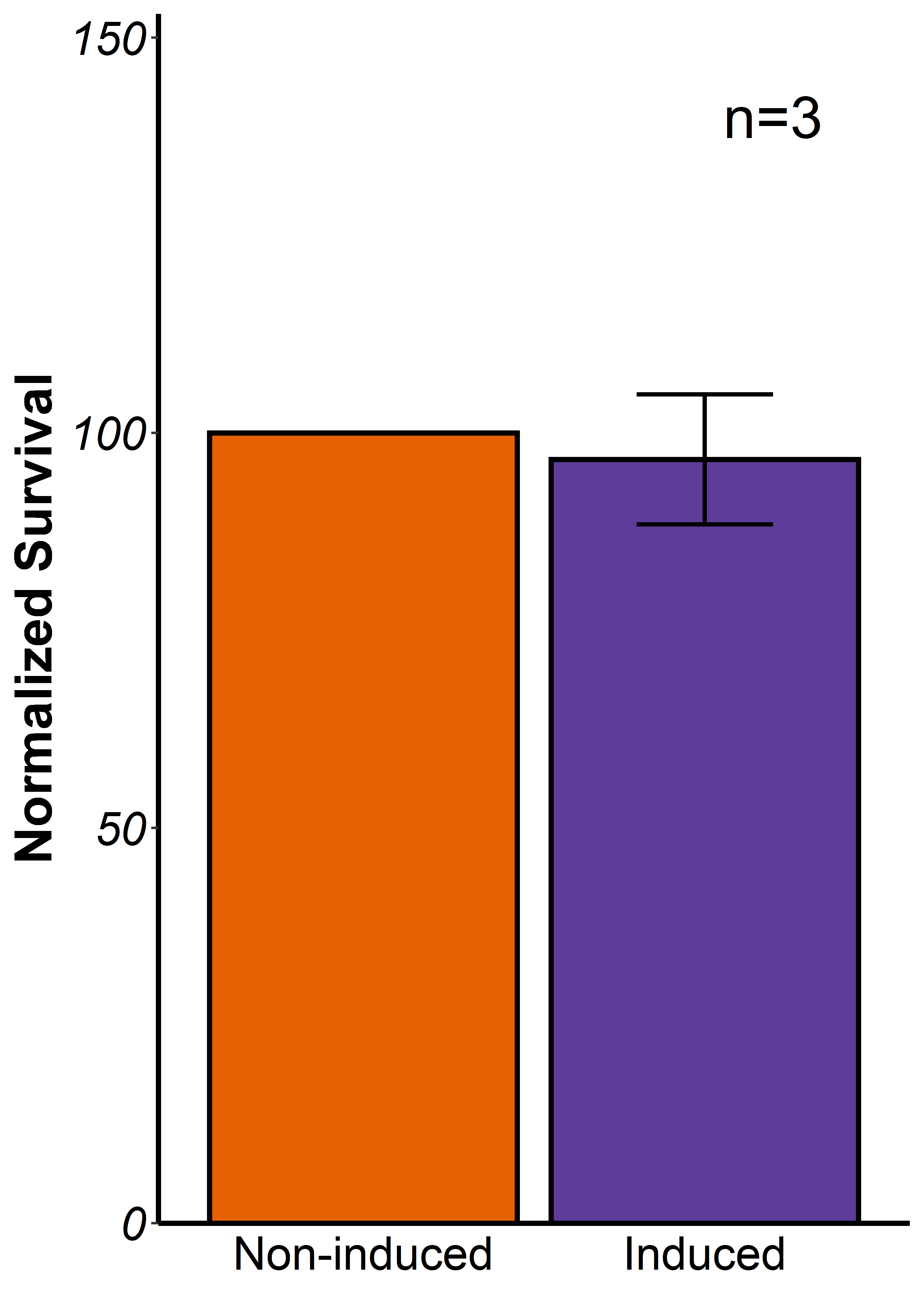

**Supplementary Figure 7 | Off-target activity assessment with the two-plasmid system.** The host strain harboring NgAgo expression plasmid and target plasmid are plated in the presence or absence of IPTG inducer and the number of colony forming units relative to the non-induced control was calculated as ‘survival’ . No significant change of survival is observed. Error bars are the standard errors generated from three replicates. Statistically significant results are indicated with * (p-value< 0.05, paired t-test).

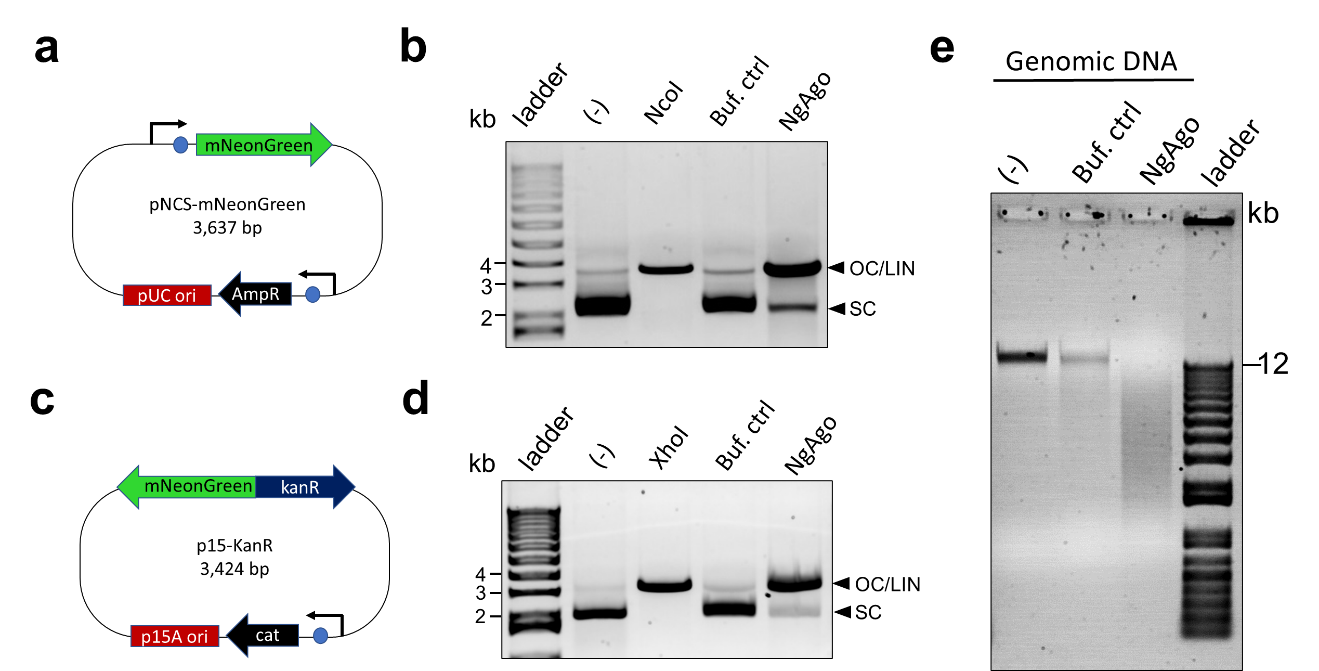

**Supplementary Figure 8 | Soluble wildtype NgAgo nicks and cuts DNA in the absence of guide DNA. a**, Plasmid map of the related plasmid, pNCS-mNeonGreen. This plasmid shares the same ampicillin antibiotic resistant gene with the NgAgo expression plasmid. **b**, Agarose gel analysis of wildtype NgAgo-treated pNCS-mNeonGreen, showing the degraded DNA product. **c**, Plasmid map of the unrelated plasmid, p15-KanR. This plasmid does not share genetic elements with the NgAgo expression plasmid. **d**, Agarose gel analysis of wildtype NgAgo-treated p15-KanR, showing the degraded DNA product. **e**, Agarose gel analysis of wildtype NgAgo-treated MG1655 genomic DNA, showing the degraded DNA product.

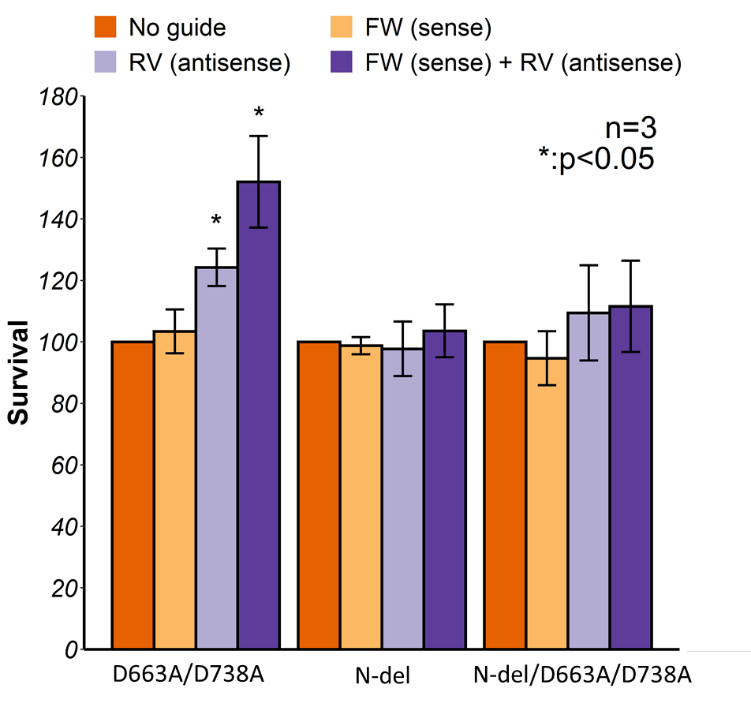

**Supplementary Figure 9 | NgAgo variants lose the ability to reduce survival.** Survival rate targeting a pseudogene (mNeonGreen) on the plasmid with NgAgo variants, including D663A/D738A, N-del, and N-del/D663A/D738A.

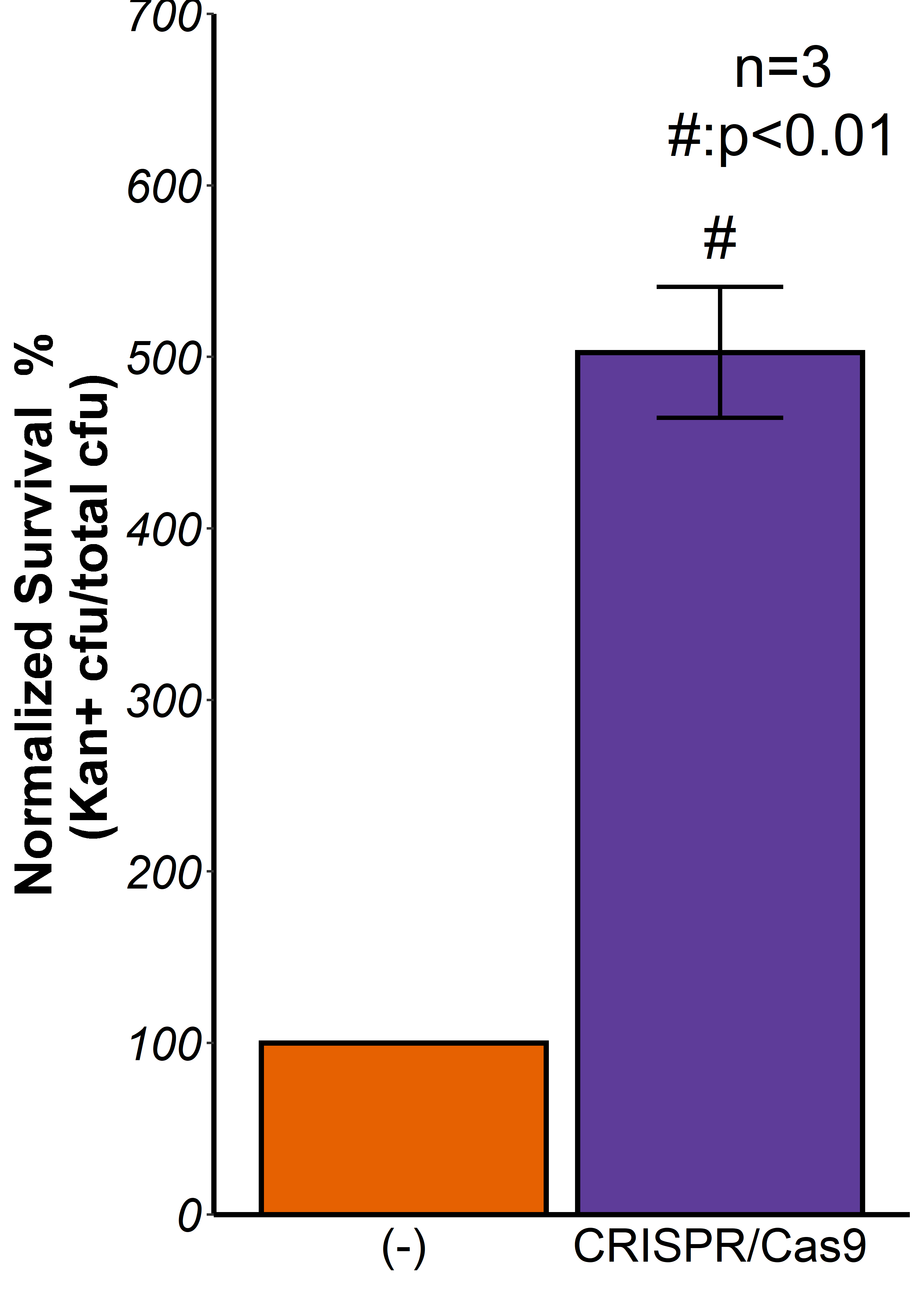

**Supplementary Figure 10 | CRISPR/Cas9 enhances homologous recombination in *E.coli*.** Induction of CRISPR/Cas9 enhance homologous recombination for 400%. Kanamycin positive colonies are normalized by total colony forming units. Error bars are the standard errors generated from three replicates. Statistically significant results are indicated with * (p-value< 0.05, paired t-test).

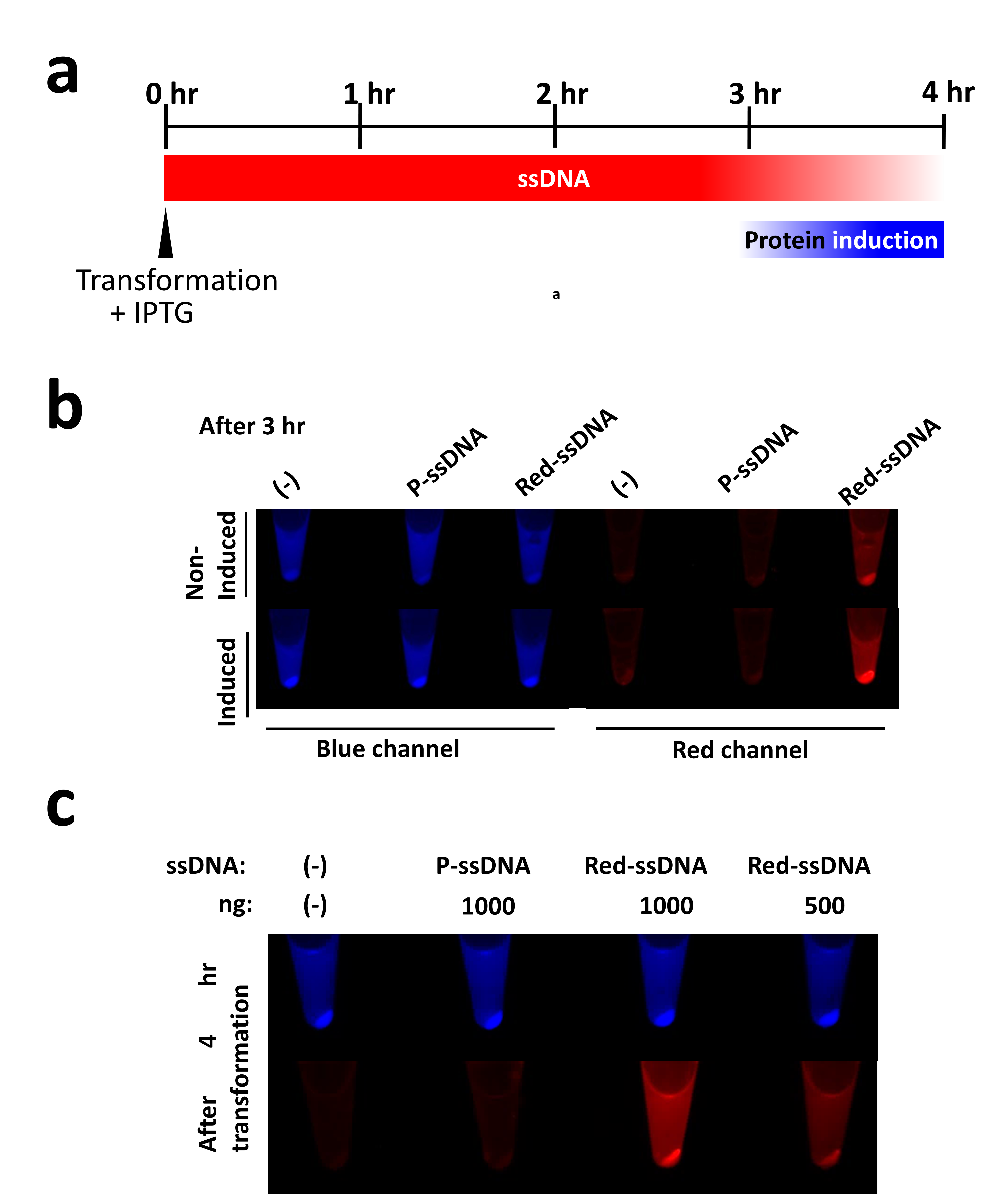

**Supplementary Figure 11 | Persistence testing of transformed guide DNA. a**, Timeline of the experimental procedure. BL21 (DE3) harboring inducible BFP expression plasmid was made electrocompetent and transformed with D4PA-labelled Red-ssDNA. After transformation, cells were resuspended in SOC in the presence of 0.1mM IPTG. **b**, BFP expression is detected after 3hr transformation. **c**, Red-ssDNA is still present after 4 hours transformation.

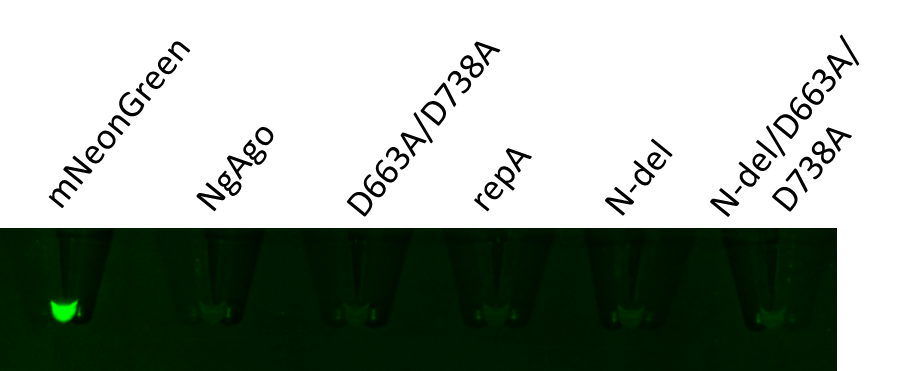

**Supplementary Figure 12** **| Production of mNeonGreen and NgAgo variants.** Cell-free system-produced mNeonGreen and NgAgo variants, including D663A/D738A, repA, N-del, and N-del/D663A/D738A, are visualized for green fluorescence to confirm successful expression in the mNeonGreen positive control.

**Supplementary Table 1.** Top 10 hits of NgAgo in Phyre 2 search.

| Structure ID | Structure source | Protein | Probability | Identity with NgAgo |
| --- | --- | --- | --- | --- |
| 5GUH | PDB | Silkworm PIWI-clade Argonaute Siwi | 100% | 15% |
| 4EI3 | PDB | Homo sapiens Argonaute2 | 100% | 18% |
| 3HO1 | PDB | Thermus thermophilus Argonaute N546 mutant | 100% | 19% |
| 4F1N | PDB | Kluyveromyces polysporus Argonaute | 100% | 14% |
| 3DLB | PDB | Thermus thermophilus Argonaute | 100% | 19% |
| 2F8S | PDB | Aquifex aeolicus Argonaute | 100% | 16% |
| 5G5T | PDB | Methanocaldcoccus janaschii Argonaute | 100% | 15% |
| 1U04 | PDB | Pyrococcus furiosus Argonaute | 100% | 12% |
| 5AWH | PDB | Rhodobacter sphaeroides Argonaute | 100% | 14% |
| 5THE | PDB | Vanderwaltozyma polyspora Argonaute | 100% | 17% |

**Supplementary Table 2.** Top 10 hits of NgAgo in HHpred search.

| Structure ID | Protein | Probability | E-value | Identity to NgAgo |
| --- | --- | --- | --- | --- |
| 5GUH | silkworm PIWI-clade Argonaute Siwi | 100% | 1e-86 | 15% |
| 4Z4D | Homo sapiens Argonaute2 | 100% | 3.4e-77 | 16% |
| [4F1N](http://www.rcsb.org/pdb/explore/explore.do?structureId=4F1N) | Kluyveromyces polysporus Argonaute | 100% | 3e-77 | 17% |
| 4NCB | Thermus thermophilus Argonaute | 100% | 2.5e-68 | 17% |
| [5G5S](http://www.rcsb.org/pdb/explore/explore.do?structureId=5G5S) | Methanocaldcoccus janaschii Argonaute | 100% | 2.6e-68 | 12% |
| 1YVU | Aquifex aeolicus Argonaute | 100% | 3.9e-68 | 16% |
| [1U04](http://www.rcsb.org/pdb/explore/explore.do?structureId=1U04) | Pyrococcus furiosus Argonaute | 100% | 1.2e-66 | 14% |
| 5I4A | Marinitoga piezophila Argonaute | 100% | 1.6e-65 | 14% |
| 6D92 | Rhodobacter sphaeroides Argonaute | 100% | 1.9e-55 | 14% |
| 5THE | Vanderwaltozyma polyspora Argonaute | 100% | 1.7e-56 | 17% |

**Supplementary Table 3**. Top 10 hits of repA domain of NgAgo in Phyre 2 search. A non-OB fold domain match was eliminated in this table.

| Structure ID | Source | Protein | Probability | Identity to NgAgo |
| --- | --- | --- | --- | --- |
| 2KEN | PDB | Methanosarcina mazei OB domain of MM0293 | 95.8% | 12% |
| 3DM3 | PDB | Methanocaldococcus jannaschii repA | 95.2% | 23% |
| 2K50 | PDB | Methanobacterium thermoautotrophicum repA-related protein | 94.6% | 12% |
| 1O7I | PDB | Sulfolobus solfataricus ssb | 94.4% | 16% |
| 1FGU | PDB | Homo sapiens REPA | 92.3% | 15% |
| d1jmca2 | SCOP | Homo sapiens RPA70 | 92% | 15% |
| 4OWX | PDB | Homo sapiens SOSS complex subunit B1 | 91.8% | 15% |
| 3E0E | PDB | Methanococcus maripaludis repA | 78.2% | 20% |
| 2K75 | PDB | Thermoplasma acidophilum OB domain of Ta0387 | 67.2% | 14% |
| d1wjja_ | SCOP | Arabidopsis thaliana hypothetical protein F20O9.120 | 66.0% | 16% |

**Supplementary Table 4**. Top 10 hits of repA domain of NgAgo in HHpred search. A non-OB fold domain match was eliminated in this table.

| Ranking | Structure ID | Protein | Probability | E-value |
| --- | --- | --- | --- | --- |
| 27 | 4OWT | Homo sapiens SOSS1 subunit B1 | 94.68% | 0.06 |
| 28 | 1WJJ | Arabidopsis thaliana hypothetical protein F20O9.120 | 94.65% | 0.086 |
| 29 | 1O7I | Sulfolobus solfataricus single-stranded DNA binding protein chain B | 94.0% | 0.28 |
| 30 | 2K50 | Methanobacterium thermoautotrophicum repA | 92.46% | 0.036 |
| 31 | [3DM3](http://www.rcsb.org/pdb/explore/explore.do?structureId=3DM3) | Methanocaldococcus jannaschii repA | 91.96% | 0.65 |
| 33 | [3E0E](http://www.rcsb.org/pdb/explore/explore.do?structureId=3E0E) | Methanococcus maripaludis repA | 88.18% | 2.5 |
| 34 | 1YNX | Saccharomyces cerevisiae repA | 87.6% | 1.3 |
| 35 | [5D8F](http://www.rcsb.org/pdb/explore/explore.do?structureId=5D8F) | Homo sapiens SOSS complex subunit B1 | 84.78% | 6.7 |
| 36 | 1JMC | Homo sapiens RPA70 | 82.12% | 4.7 |
| 37 | [4HIK](http://www.rcsb.org/pdb/explore/explore.do?structureId=4HIK) | Schizosaccharomyces pombe Pot1pC | 81.44% | 5.1 |

**Supplementary Table 5.** DNA primers used in this study. Restriction enzyme recognition sites are underlined and indicated in the primer name.

| **Name** | **Sequences (5’>3’)** | **Template** | **Used to construct** |
| --- | --- | --- | --- |
| 5' NcoI 3xG Ago | ATCACCATGGGTGGCGGTATGGTGCCAAAAAAGAAGAG | Nls-NgAgo-GK | pET-GST-Ago-His |
| 3' XhoI Ago | ATCACTCGAGCTTACTTACATATGGATCCCGG |  |  |
| NdeI HIS-Ago 5 | TATACATATGGGTCACCATCATCATCACCATTCATCGCATCACCATCACCATCACGTGCCAAAAAAGAAGAG | Nls-NgAgo-GK | pET-His-Ago |
| XhoI rmNdeI Ago 3' | ATATCTCGAGTTACTTACTTACGTATGGATCCCGG | Nls-NgAgo-GK | pET-His-Ago |
| XhoI STOP repA 3' | CTAACTCGAGTTACTCGACGGTCGTCTGG | Nls-NgAgo-GK | pET-His-repA |
| D663A 3' | CGGGGTAGCTCCGAGAGACCGCAATCCCAATGAACATATC | pET-His-Ago | NgAgo mutant |
| D663A 5' | GATATGTTCATTGGGATTGCGGTCTCTCGGAGCTACCCCG | pET-His-Ago |  |
| D738A 5' | CGACCCATATCGTCATCCACCGTGCGGGCTTCATGAACGAAGACCTCGAC | pET-His-Ago | NgAgo mutant |
| D738A 3' | GTCGAGGTCTTCGTTCATGAAGCCCGCACGGTGGATGACGATATGGGTCG | pET-His-Ago |  |
| XbaI KanR 5' | ATGGTCTAGAATGGGATCGGCCATTG | pTKIP-neo | kanR-mNeonGreen cassette integration |
| BamHI KanR 3' | ATTTGGATCCTTAGAAGAACTCGTCAAGAAGGC | pTKIP-neo |  |
| XbaI tGreen 5' | CCATTCTAGACCATGGTAGATGGCTCCG | pNCS-mNeonGreen |  |
| XhoI Green uni 3' | TGATCTCGAGAGAGAATATAAAAAGCCAGATTATTAATCCGGCTTTTTTATTATTTTTACTTGTACAGCTCGTCCATGC | pNCS-mNeonGreen |  |
| XbaI tKanR 5' | TAGCTCTAGAGAAAGAGGAGAAATACTAGATGGGATCGGCCATTG | pTKIP-neo | donor plasmid p15-KanR-PtetRed |
| EcoRI tKanR 3' | ATATGAATTCGATACTTTCTCGGCAGGAGC | pTKIP-neo |  |
| XbaI J23100 tGreen 5' | TTTCTCTAGAGCTAGCACTGTACCTAGGACTGAGCTAGCCGTCAACCATGGGAAGCCACATC | pNCS-mNeonGreen |  |
| XhoI Green uni 3' | TGATCTCGAGAGAGAATATAAAAAGCCAGATTATTAATCCGGCTTTTTTATTATTTTTACTTGTACAGCTCGTCCATGC | pNCS-mNeonGreen |  |
| XhoI pTet 5' | ATCACTCGAGTCCCTATCAGTGATAGAGATTGACATCCCTATCAGTGATAGAGATACTGAGCACTCTAG | pTK-Red |  |
| XhoI DT exo 3' | TGATCTCGAGAAAAAAAAACCCCGCCGAAGCGGGGTTTTTTTTTTCATCGCCATTGCTCC | pTK-Red |  |

**Supplementary Table 6.** DNA guides used in this study.

| **Name of 5’ phosphorylated guide DNA** | **Sequences (5’ to 3’)** |
| --- | --- |
| FW p-tetA | P-GGATTGGCCTTATCATGCCAGTCT |
| RV p-tetA | P-AGACTGGCATGATAAGGCCAATCC |
| FW p-mNeonGreen | P-TTAACTACCGCTACACCTACGAGG |
| RV p-mNeonGreen | P-CCTCGTAGGTGTAGCGGTAGTTAA |
| FW p-arpB | P- ATACAGCAGCATGTCCCCTTAGTC |
| RV p-arpB | P- GACTAAGGGGACATGCTGCTGTAT |
| D4PA-labelled Red-ssDNA | D4PA-TTAACTACCGCTACACCTACGAGG |

**WORK CITED**

1.	Swarts, D. C. *et al.* Autonomous Generation and Loading of DNA Guides by Bacterial Argonaute. *Molecular Cell* **65**, 985-998.e6 (2017).

2.	Zander, A. *et al.* Guide-independent DNA cleavage by archaeal Argonaute from Methanocaldococcus jannaschii. *Nature Microbiology* **2**, 17034 (2017).

3.	Hunt, E. A., Evans Jr, T. C. & Tanner, N. A. Single-stranded binding proteins and helicase enhance the activity of prokaryotic argonautes in vitro. *PloS One* **13**, e0203073 (2018).

4.	Niu, Y., Tenney, K., Li, H. & Gimble, F. S. Engineering variants of the I-SceI homing endonuclease with strand-specific and site-specific DNA-nicking activity. *Journal of Molecular Biology* **382**, 188–202 (2008).
